# Beyond the Sin3/HDAC Complex: FAM60A emerges as a regulator of RNA Splicing

**DOI:** 10.1101/2024.01.16.575949

**Authors:** Md Nazmul Huda, Rosalyn Zimmermann, Preeti Nagar, Jahangir Alam, Md Rafikul Islam, Dayebgadoh Gerald, Mojnu Miah, Cassandra G Kempf, Janet L. Thornton, Ying Zhang, Zhihui Wen, Gaye Hatem, Allen Gies, Stephanie Byrum, Mohammad Alinoor Rahman, Laurence Florens, Michael P. Washburn, Sayem Miah

## Abstract

FAM60A, traditionally linked to chromatin remodeling within the Sin3/HDAC complex, has emerged as a critical regulator in RNA splicing. Employing an integrative approach that combines immunological assays, CRISPR/Cas9 technology, comprehensive genomics, proteomics, and advanced cross-linking mass spectrometry, complemented by sophisticated 3D molecular modeling, our study challenges and extends the existing understanding of FAM60A’s functional dynamics. Contravening previous perceptions, our findings elucidate that FAM60A does not interact directly with SIN3A, rather establishes direct interactions with SAP30 and HDAC1, redefining its relationship with the Sin3/HDAC complex. These interactions, deciphered through detailed 3D structural analysis supported by cross-linking constraints, signify a complex architectural role of FAM60A within chromatin remodeling processes. Moreover, our research unveils FAM60A’s pivotal role in RNA processing, particularly in splicing regulation. Through extensive molecular interactions with a diverse array of mRNA-binding proteins and principal spliceosome components, FAM60A emerges as a key regulator of RNA splicing. This expanded role delineates its influence on gene expression regulation, spotlighting its capacity to modulate critical cellular processes. In sum, this study unveils FAM60A’s key role in gene regulation and RNA splicing, and suggests new paths for cellular and therapeutic research.

## Introduction

Chromatin remodeling complexes, such as the Sin3/Histone Deacetylase (HDAC), are essential in regulating the dynamic structure of chromatin. This modulation is critical for various DNA-dependent processes, including DNA repair, replication, and transcription^1–4^. The Sin3/HDAC complex, integral to chromatin modification, plays a significant role in gene regulation by facilitating histone deacetylation. This deacetylation compacts chromatin into a heterochromatin state, subsequently suppressing gene transcription^2,5^. Within this multi-subunit complex, the FAM60A subunit^6^ is notable for its role in directing the complex’s specific interactions with diverse genomic regions^7^. Despite its established role within the Sin3/HDAC complex, the cellular functions of FAM60A beyond this complex are still largely unexplored.

In proteomic studies, FAM60A has been identified as a key subunit of the Sin3/HDAC complex, primarily involved in gene suppression within the TGF-beta signaling pathway^6^. This metazoan-specific protein is exclusive to the Sin3 complex and does not interact with other complexes^6^. Its significance is further highlighted by its variable expression in U2OS cell cycles^8^ and its role in embryonic stem cells, particularly through interactions with SIN3A^7,9^. FAM60A’s widespread presence across various mammalian cell types and its pivotal role in modulating TGF-beta signaling underscore its importance in chromatin remodeling and gene regulation, confirming its significance in these pathways.

Additionally, FAM60A is highly expressed in embryonic stem cells^7^ and essential for normal mouse development^9^. The absence of FAM60A results in underdeveloped organs and embryonic fatality, significantly affecting DNA methylation at gene promoters^9^. In cancer research, however, FAM60A’s role is complex and dualistic. While some studies categorize FAM60A as a tumor suppressor^6^, other research links it to tumor progression and drug resistance^10–12^. This paradoxical nature underscores the complexity of FAM60A’s function in cellular biology and highlights the need for further investigation to fully understand its role in biological processes.

This study focuses on exploring the diverse roles of FAM60A beyond its interaction with the Sin3/HDAC complex. We used a combination of immunological, biochemical, and CRISPR/Cas9 techniques, along with genomics, proteomics, and chemical cross-linking mass spectrometry (XL-MS), supported by computer-based molecular modeling. Our findings show that FAM60A directly interacts with the HDAC1 and SAP30 subunits, connecting it to SIN3A. Importantly, we also discovered FAM60A’s involvement in RNA binding and its crucial role in regulating RNA splicing. This research not only deepens our understanding of FAM60A as an important RNA-binding protein but also sheds light on its key role in RNA splicing, a vital process in cell biology.

## Results

### FAM60A is an integral component of the Sin3/DHAC histone deacetylase complex

Earlier research highlighted FAM60A co-purified with several Sin3/HDAC complex members, including BRMS1/1L, SAP30/30L, and ING1/2^6,13–15^. To elucidate the interaction network among these proteins and discern relationships between the core subunits of the Sin3/HDAC complex and FAM60A, we integrated the Halo affinity tag with Multidimensional Protein Identification Technology (MudPIT). This approach (**Fig. 1A**), was validated by expressing Halo-SIN3A in HEK 293 cells, purifying associated protein complexes, and quantifying these complexes using MudPIT mass spectrometry and visualized using Cytoscape^16^ (**Fig. 1B** and **Suppl. Table S1**). Notably, endogenous FAM60A co-purified with Halo-SIN3A alongside other Sin3/HDAC complex subunits such as HDAC1, HDAC2, SUDS3, SAP30, SAP30L, BRMS1, ING1, and ING2 (**Fig. 1B**, highlighted in pink). The interaction of Sin3/HDAC complex subunits with SIN3A was further corroborated through immunoblotting. After expressing Halo-SIN3A in HEK 293 cells, we affinity-purified SIN3A-associated proteins, performed SDS PAGE-western blotting, and probed them with antibodies targeting core Sin3/HDAC complex subunits, including FAM60A (**Fig. 1C**). To confirm FAM60A as a core component of SIN3A-containing complexes, we ectopically expressed Halo-tagged versions of core components of the Sin3/HDAC complex, including SIN3A, HDAC1, HDAC2, SUDS3, SAP30, SAP30L, ARID4A, BRMS1, ING1, and ING2. Subsequent affinity purification and quantification of these complexes via MudPIT mass spectrometry are presented in a heatmap. (**Fig. 1D** and **Suppl. Table S1**). We observed that endogenous FAM60A co-purified with all core subunits of this complex, except ING1, ARID4A, and HDAC2. It is to note that ARID4A, a transcription factor co-purifying with SIN3A, is not traditionally considered a Sin3/HDAC complex subunit.

**Fig. 1:**
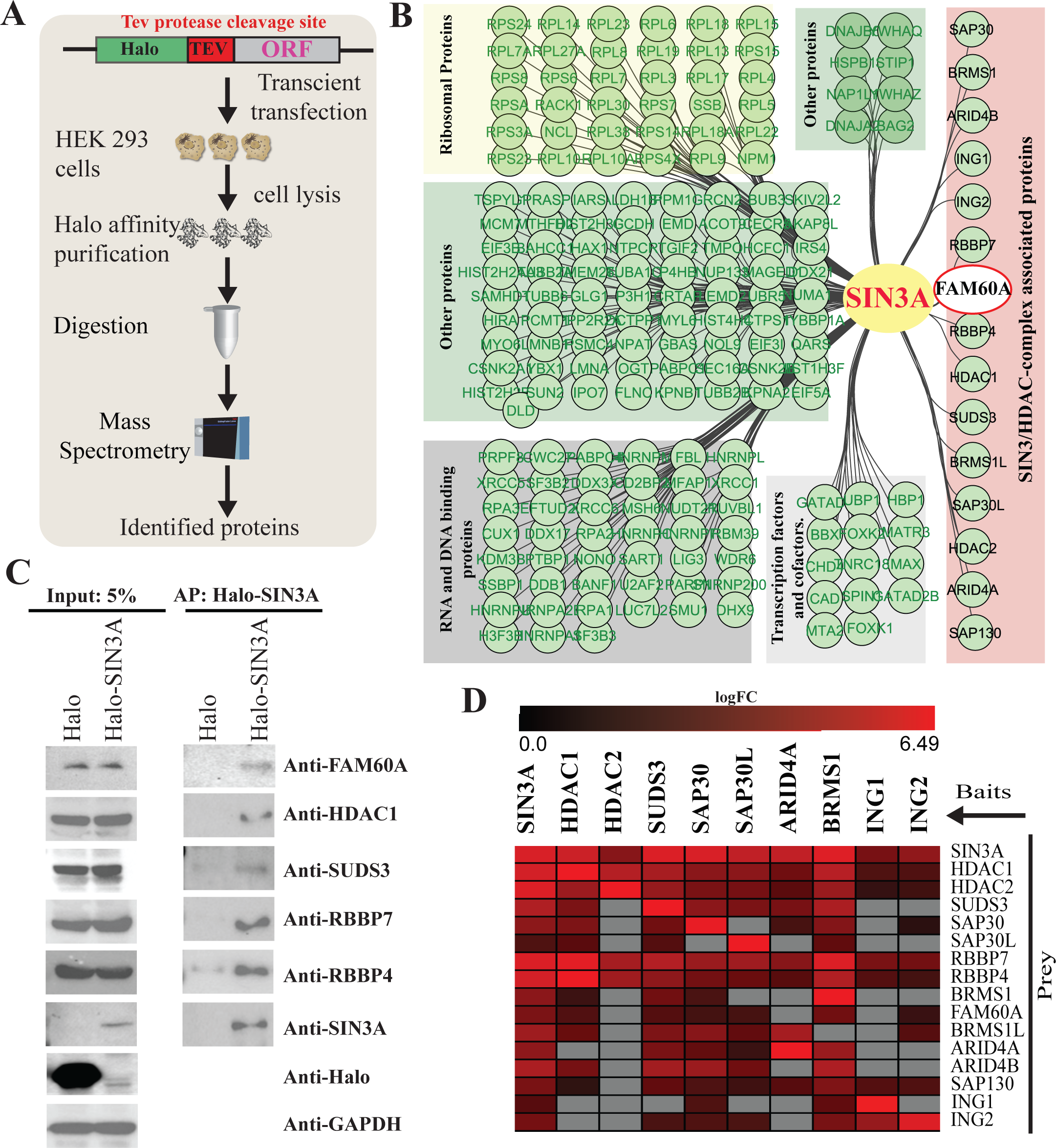
SIN3A network reveals FAM60A as a subunit of Sin3/HDAC. a, a schematic of refined proteomics workflow employing Mass spectrometry for protein identification. b, HEK293T cells were transfected with Halo-SIN3A and the cell lysates were subjected to affinity purification with Halo magnetic beads. Purified protein complexes were identified by Mass-spec and analyzed by QSPEC (QSPEC log2FC ≥ 1, QSPEC FDR < 0.05) and visualized in a differential network using Cytoscape. c, Halo-SIN3A or Halo plasmid alone was ectopically expressed in HEK293T cells followed by Halo affinity purification. The Halo affinity purified protein were subjected to immunoblotting with anti-FAM60A, anti-HDAC1, anti-SUDS3, anti-RBBP7, anti-RBBP4, anti-SIN3A, and anti-SMAD4 antibodies. Total cell lysates were also analyzed by immunoblotting using same antibodies and anti-GAPDH as a loading control. d, identified subunits of Sin3/HDAC complex were N-terminally Halo tagged and ectopically expressed in HEK 293 cells followed by affinity purification and identified using Mass-spectrometry analysis (QSPEC log2FC ≥ 1, FDR < 0.05) and visualized in a heatmap.

Next, to determine what portion of SIN3A might interact with FAM60A, we generated a series of deletion mutants. Since SIN3A is a scaffolding protein, we systematically deleted highly conserved regions annotated as specific domains **(Fig. S1A)**: four Paired Amphipathic Helices, (PAH1-4); SUDS3 and SAP130 interacting domain; HDAC Interacting Domain (HID); Highly Conserved Region, (HCR) at the C-terminus; 7 poly glutamic acids domain; and Proline-rich domain, however, SIN3A lacks enzymatic activities and DNA binding motifs^5^. All nine-deletion mutants of SIN3A were Halo-tagged and expressed in HEK 293 cells and probed with anti-Halo antibody **(Fig. S1B).** We also checked whether the intracellular localization of those SIN3A deletion mutants were altered due to the lack of conserved domains. We used live-cell imaging to assess the localization of ectopically expressed Halo-SIN3A and Halo-SIN3A deletion mutants by transfecting HEK293T cells and imaging by confocal microscopy **(Fig. S1C)**. We observed that Halo-SIN3A and all deletion mutants predominantly localized to the nucleus, while Halo-SIN3A_Δ8 that lacks the 7 poly-Glu domain mis-localized to the cytosol **(Fig. S1C)**.

To identify the essential domains in SIN3A for FAM60A binding, we conducted advanced proteomic analysis on Halo-SIN3A deletion mutants. Halo-tagged SIN3A deletion mutants were ectopically expressed, purified using Halo affinity purification, and then analyzed with liquid chromatography-mass spectrometry (LC-MS) to identify proteins interacting with the deletion mutants. The identified interacting proteins of each Halo-SIN3A deletion mutants are presented in a heatmap **(Fig. S1D** and **Suppl. Table S1)**. We observed that SIN3A and its deletion mutants interact with hundreds of proteins including the core subunit of Sin3/HDAC complex **(Fig. S1D).** We then focused on the core subunit of Sin3/HDAC complex and found that FAM60A did not interact with SIN3A lacking HCR (SIN3A_Δ7_968-1186) along with BRMS1 and ING1**(Fig. S1E** and **Suppl. Table S1)**. However, unlike FAM60A, BRMS1 did not interact with SIN3A_Δ6, SIN3A_Δ7, SIN3A_Δ8, while ING1 did not interact with SIN3A_Δ4, SIN3A_Δ5, SIN3A_Δ6, SIN3A_Δ7, and SIN3A_Δ8. We also noticed that SUDS3 did not interact with SIN3A_Δ5 and SAP130 did not interact with SIN3A_Δ3 mutants. We also identified domain-specific interaction of SIN3A with different transcription factors such as ARID4A, ARID4B, and MAX **(Fig. S1E)**. Overall, our data suggest that FAM60A is a core component of the Sin3/HDAC complex and binds to the C-terminal HCR region of SIN3A.

### Diverse protein interaction network of FAM60A extends beyond the Sin3/HDAC complex

FAM60A is a highly conserved protein across many vertebrate **(Fig. S2A)** suggesting its important role in physiological processes. To uncover FAM60A’s biological functions, we generated a series of Halo-tagged FAM60A deletion mutants (**Fig. 2A**). To assess the proper expression and localization of Halo-FAM60A deletion mutants, we overexpressed them in HEK293T cells and visualized their localization using live-cell imaging (**Fig. 2B**). Additionally, we performed Halo affinity purification followed by gel electrophoresis and silver staining to visualize the affinity-purified proteins **(Fig. S2B)**. Subcellular localization analysis revealed that the full-length Halo-FAM60A, along with its deletion mutants FAM60A_Δ1 (consisting of amino acids (AAs) 1-148), FAM60A_Δ2 (only containing the GATA like zinc finger domain, AAs 1-95), and the spliced isoform FAM60A_Δ4 (only containing the C-terminal AAs 149-221), were localized to the nucleus, while mutant FAM60A_Δ3 (consisting of AAs 96-148 only) exhibited non-nuclear localization (**Fig. 2B**). This observation aligns with our silver staining analysis, where FAM60A_Δ3 did not interact with other proteins **(Fig. S2B)**, suggesting potential instability or improper expression of this truncated protein fragment.

**Fig. 2:**
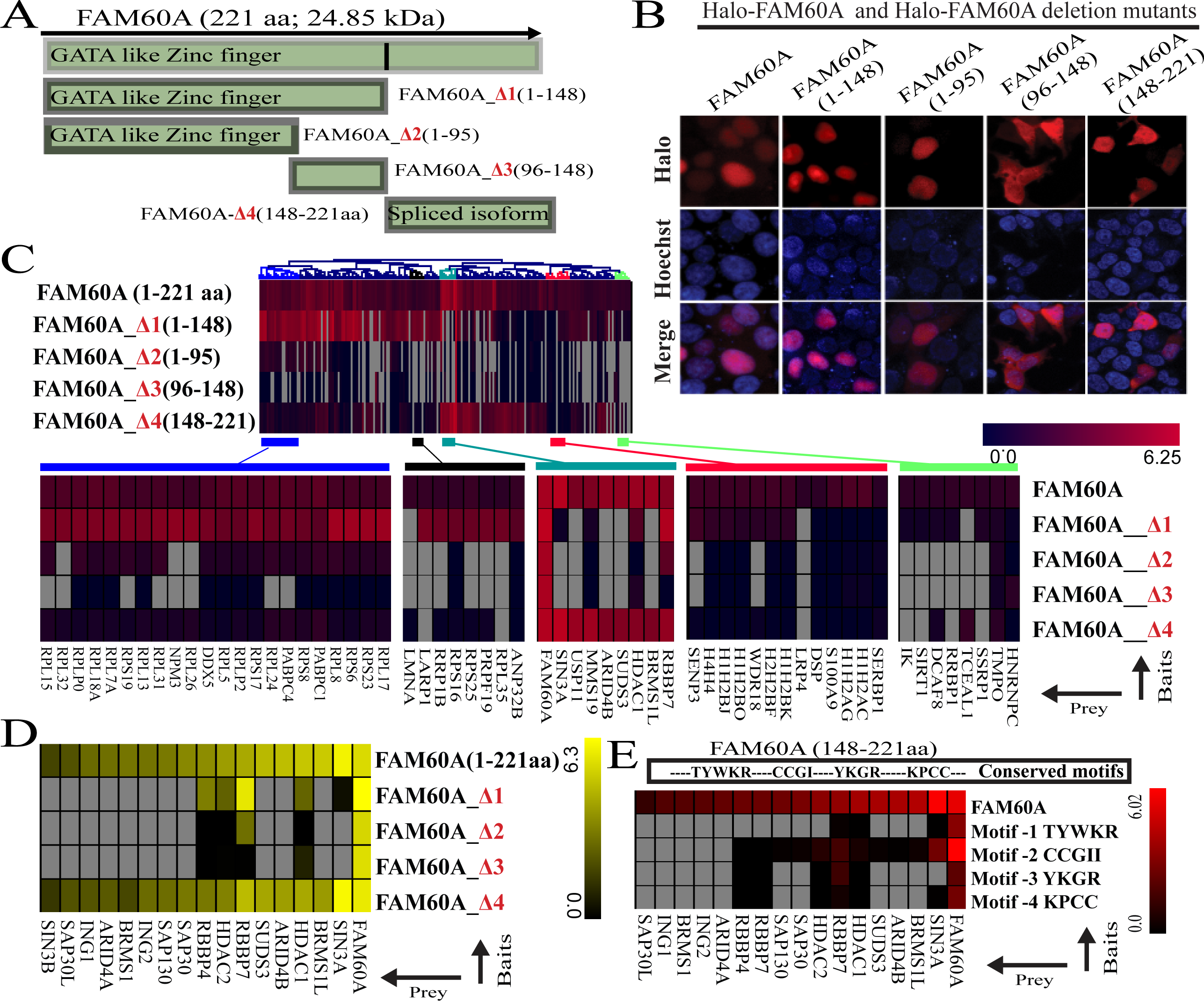
Enhanced deletion analysis of the FAM60A Protein. a, schematic-diagram illustrating various deletions in the *FAM60A* gene. b, Halo-tagged FAM60A deletion mutants were transfected into HEK293T cells, labeled with Halo-Tag TMRDirect fluorescent ligand (red), and DNA was stained using Hoechst dye (blue). c, Halo-FAM60A deletion mutants were expressed in HEK293T cells, purified, and analyzed via proteomics (QSPEC log2FC ≥ 1, FDR < 0.05). Results were displayed in a heatmap (Red, black). d, the interactions of full length and truncated FAM60A with Sin3/HDAC complex core-subunits were visualized in a heatmap (grey indicate no interaction). e, heatmap showing FAM60A conserved motif-specific interactions with Sin3/HDAC complex core-subunits (Grey Signify No Interaction).

Next, FAM60A and its mutants were affinity purified and analyzed by proteomics. We observed that FAM60A and its four mutants interacted with hundreds of proteins, which are presented in a heatmap using Genesis^17,18^ (**Fig. 2C and Suppl. Table S2**). To elucidate FAM60A’s protein interactions and their functional implications, we examined its interaction network, as depicted in **Fig. S3A**. This analysis uncovered different interaction patterns for full-length FAM60A and its variants FAM60A_Δ1 and FAM60A_Δ4. Most components of the Sin3/HDAC complex were found to interact with both full-length FAM60A and FAM60A_Δ4 (the C-terminus of FAM60A from AAs 149-221). In contrast, FAM60A_Δ1 (FAM60A N-terminal from AAs 1-148 including the GATA like zinc finger domain) predominantly interacted with RNA-binding proteins, including NPM1 and NCL, suggesting a role in RNA processing beyond chromatin remodeling. Confirmatory immunoblotting analysis with core Sin3/HDAC complex subunits and FAM60A aligned with our AP-MS results (**Fig. S3B**).

FAM60A predominantly interacts with the core elements of the Sin3/HDAC complex via its C-terminus. Yet, the submodule module consisting of HDAC1, HDAC2, RBBP4, and RBBP7 interacted with both the N- and C-terminus regions of FAM60A (**Fig. 2D**, **Suppl. Table S2 and Fig. S3A**). Since FAM60A predominantly interacts with the Sin3/HDAC complex via its C-terminus, we examined four conserved amino acid motifs located in the C-terminus of FAM60A to better dissect this interaction (**Fig. S3C**). The motifs in FAM60A were meticulously altered by substituting amino acids with structurally similar counterparts, such as replacing tyrosine (Y) with phenylalanine (F), to minimize the impact on structure and function while preserving the protein’s overall integrity. These modified versions of FAM60A were then introduced into HEK293 cells. Following affinity purification and proteomic analysis, it became evident that Motif-1 (TYWKR) is crucial for FAM60A interaction with the Sin3/HDAC complex, while Motif-3 and 4 also play significant roles. However, Motif-2 did not appear to be important in this interaction (**Fig. 2E** and **Suppl. Table S2**).

### Size exclusion chromatography and XL-MS reveal core structure of FAM60A-Sin3/HDAC complex

To address the challenges in identifying the distinct multi-subunit protein complexes and their core components that may interact with FAM60A, we further fractionated the eluate obtained from the Halo-FAM60A affinity purification by size exclusion chromatography (**Fig. 3A**). We specifically processed the affinity-purified FAM60A by loading it onto a Superose 6 column and collecting 48 fractions of 500μl each. From these, 20 fractions (14-33) were selected, digested, and then analyzed using MudPIT. Although FAM60A’s expected molecular weight is 29kDa, its fractionation profile confirmed it assembles into larger complexes. To elucidate the molecular composition of potential FAM60A-Sin3/HDAC complexes, we assessed whether components of this complex were present. Our observations revealed that FAM60A co-fractionated with SIN3A, RBBP7, RBBP4, HDAC1, HDAC2, and SUDS3 (**Fig. 3B** and **Suppl. Table S3**).

**Fig. 3:**
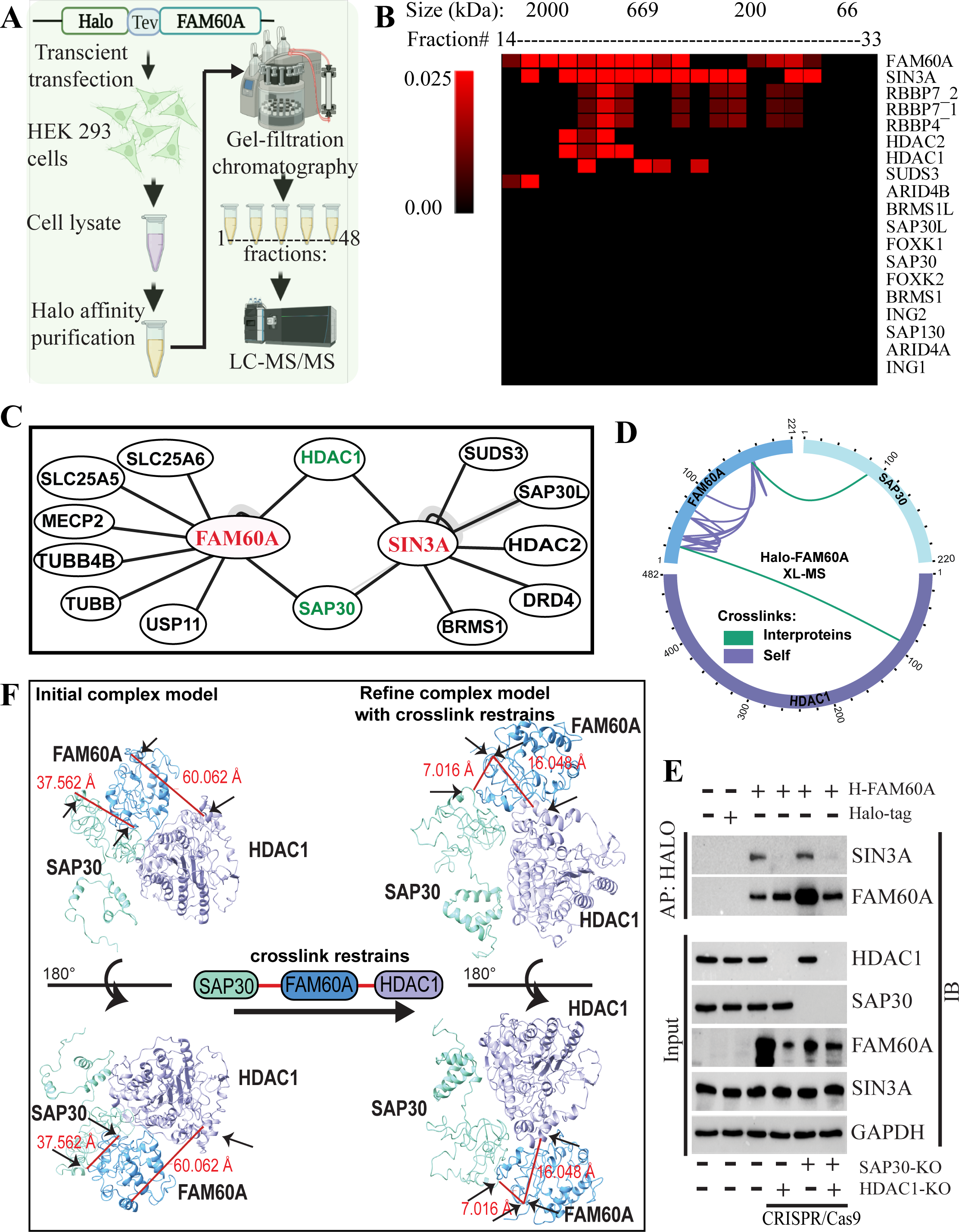
Architecture of the FAM60A-SIN3A/HDAC1 Complex. a, workflow for Halo-FAM60A-expressing HEK 293 cell extract preparation, HaloTag purification, size exclusion chromatography, and AP-MS analysis. b, Halo-FAM60A affinity purified proteins were fractionated through size-exclusion chromatography and analyzed by LC-MS and visualized in a heatmap (Red, and Black). c, Halo-FAM60A affinity purified samples were treated with DSSO followed by high-resolution Mass spectrometry analysis. High-resolution MS analysis identifies the FAM60A-HDAC1 and FAM60A-SAP30 crosslinked peptide. d, crosslink maps for FAM60A-HDAC1 and FAM60A-SAP30: These maps display the cross-links involving proteins identified in our analysis, which were enriched by Halo-FAM60A. e. *HDAC1* and *SAP30* were individually or simultaneously knocked out with CRISPR/Cas9. Subsequently, Halo-FAM60A was overexpressed, and protein complexes were purified through affinity purification. These complexes were then subjected to immunoblotting analysis and probed with various antibodies. f, structures of FAM60A were modeled using HADDOCK, guided by docking restraints derived from the indicated crosslinks, which facilitated the generation of an initial complex model. Further refinement resulted in a model of the FAM60A/HDAC1/SAP30/SIN3A sub-structure, demonstrating the FAM60A residues crosslinked to HDAC1 and SAP30, and elucidating the crosslinking of HDAC1 and SAP30, which connect them to SIN3A.

After mapping the interaction network of FAM60A (**Fig. 2**), and its co-fractionated partners (**Fig. 3B**), we aimed to understand the direct interactions within this network and how FAM60A influences these interactions. Using a crosslinking mass spectrometry (XL-MS) approach^15,19^, we treated the eluate from Halo-FAM60 affinity purification with the mass spectrometry cleavable crosslinker disuccinimidyl sulfoxide (DSSO) before LC/MS analysis. The detected crosslinked peptides included pairs of residues within the FAM60A-Sin3/HDAC complex that are likely in close proximity, less than 30Å apart (**Fig. 3C– 3D** and **Suppl. Table S3**). We discovered that FAM60A and SIN3A cross-linked with SAP30 and HDAC1, yet no cross-links were observed between FAM60A and SIN3A (**Fig. 3C–3D**). Additionally, we also identified 13 self-crosslinks of FAM60A as shown in **Fig. 3D**.

To corroborate whether the interaction between FAM60A and SIN3A, was mediated by SAP30 and HDAC1, we performed targeted knockouts of SAP30 and HDAC1, both individually and in combination, in HEK293 cells (**Fig. 3E**). Post-knockout, we ectopically expressed Halo-FAM60A, followed by affinity purification and immunoblotting analysis with a SIN3A-specific antibody. Our results showed that the FAM60A-SIN3A interaction was disrupted in HDAC1 knockout cells, but remained unaffected in SAP30 knockout cells (**Fig. 3E**). Furthermore, given the observed co-fractionation of SUDS3 with FAM60A (**Fig. 3B**), we examined the FAM60A-SIN3A interaction in the background of SUDS3 knockout, and combined SUDS3 and HDAC1 knockouts. We observed a notable reduction in the FAM60A-SIN3A interaction in both HDAC1 knockout and SUDS3-HDAC1 double knockout cells, while this interaction persisted in cells with SUDS3 knockout (**Fig. S4A**).

Next, we used the crosslinking data to guide the docking of FAM60A structure with its direct interaction partners, to unravel the three-dimensional architecture of the FAM60A complex. Our analysis revealed significant crosslinks between FAM60A and two members of the Sin3/HDAC complex: HDAC1 (between amino acids FAM60AK22 and HDAC1K90) and SAP30 (between amino acids FAM60AK168 and SAP30K87) (**Fig. 3D** and **Suppl. Table S3**). To enhance our understanding of the potential structural organization of these proteins within a cellular context, we predicted protein structures using advanced computational tools such as Alphafold^20^, I-TASSER, and I-TASSER MTD^21^. Utilizing the HADDOCK platform^22^, we guided the docking of FAM60A, HDAC1, and SAP30 structures. Our model structure was further refined by incorporating both interprotein and self-crosslinks of FAM60A as docking restraints. This comprehensive approach led to a refined model structure of the FAM60A, HDAC1, and SAP30 complex (**Fig. 3F**), providing a clearer picture of the intricate interactions within the Sin3/HDAC complex.

### FAM60A exhibits an affinity for RNA-binding proteins and has the ability to bind with mRNA

Besides binding with the components of the Sin3/HDAC complex, Halo-FAM60A interacts with a large number of RNA binding proteins (**Fig. 2C** and **Fig. S3A**). To minimize potential effects from the Halo-tag at FAM60A N-terminus, we moved the tag to the C-terminus and expressed it in HEK293 cells. No interference in nuclear localization was observed (**Fig. 4A**). We then purified FAM60A-Halo containing complexes and identified a set of 178 proteins consistently copurifying with both tagged versions of FAM60A (**Fig. 4B** and **Suppl. Table S4**). We next analyzed the gene ontology (GO) terms associated with this set of 178 FAM60A associated proteins. In addition to enrichment for GO terms related to histone deacetylase complexes (**Fig4B** – Sin3-type complex), we were surprised to find an enrichment for a set of GO terms related to intracellular ribonucleoprotein complex, spliceosomal snRNP complex, spliceosomal complex and mRNA stability complex (**Fig. 4B**). We anticipated that a proportion of the ribosomal proteins might be associated with FAM60A indirectly via interactions with other proteins or RNA. We sought to identify the most abundant FAM60A associated proteins with the associated GO term “intracellular ribonucleoprotein complex” and we found that NPM1 and a second nucleolar protein, NCL, also involved in ribosome synthesis, were the two most abundant of this group of proteins, and RNA processing proteins in both Halo-FAM60A and FAM60A-Halo purifications (**Fig. 4C** and **Suppl. Table S4**). Remarkably, these analyses highlighted FAM60A strong associations with key RNA processing assemblies, including the spliceosome, mRNA stability, snRNP, and ribonucleoprotein complexes, underscoring its pivotal role in RNA processing.

**Fig. 4:**
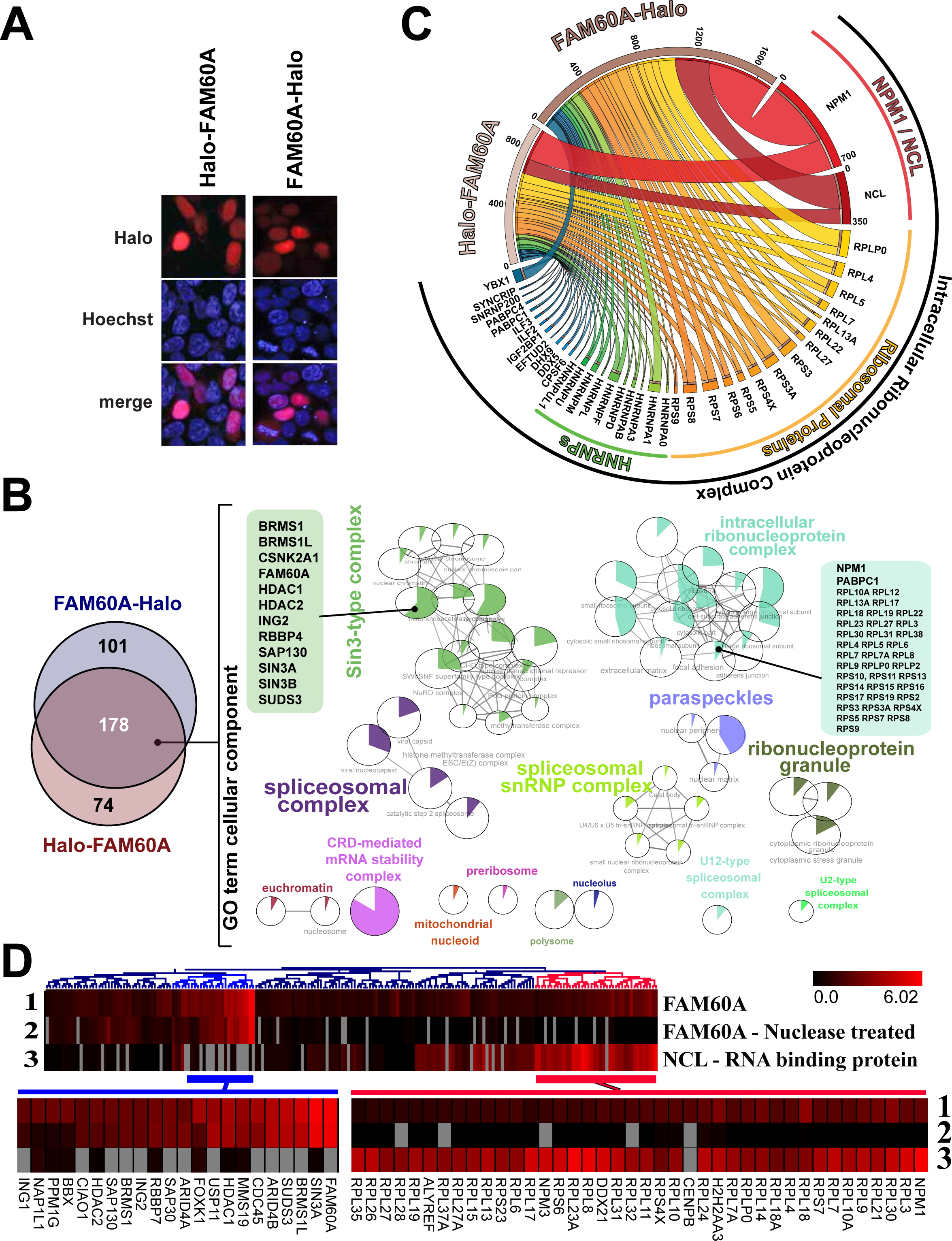
FAM60A interaction with RNA processing proteins. a, Halo-FAM60A or FAM60A-Halo was transfected into HEK 293 cells, labeled with Halo-Tag TMRDirect fluorescent ligand (red), and DNA was stained using Hoechst dye (blue). b, proteins associated with Halo-FAM60A and FAM60A-Halo were subjected for proteomic analysis (with a Z score ≥ 3 and FDR ≤ 0.05) and were evaluated for their cellular components using the ClueGo plugin in Cytoscape, with P values ≤ 0.05. c, Halo-FAM60A or FAM60A-Halo expressing HEK293 cells were subject to Halo affinity purification and analyzed by LC-MS and visualized in a CIRCOS plot. d, HEK293 cells were transfected with Halo-FAM60A, which then underwent an affinity purification process followed by nuclease treatment. Both nuclease-treated and untreated proteins underwent proteomics analysis (QSPEC log2FC ≥ 1, FDR < 0.05), visualized in a heatmap. Halo-NCL was a positive control for RNA binding and processing protein.

To understand whether FAM60A directly interacts with RNA binding proteins, we ectopically expressed Halo-FAM60A and affinity-purified FAM60A-associated proteins. Prior to proteomic analysis, we treated these proteins with Benzonase®, an endonuclease that digests both DNA and RNA, to eliminate nucleic acid-mediated interactions of FAM60A with RNA binding proteins. Proteomic analysis of affinity-purified FAM60A proteins demonstrated its interaction with numerous RNA and ribosomal RNA binding proteins (**Fig. 4D** and **Suppl. Table S4**). Notably, the number of RNA interacting partners of FAM60A was comparable to NCL, a known RNA-binding protein^23,24^, which served as a positive control for this study. Intriguingly, most nucleic acid-mediated interactions were diminished after nuclease treatment, suggesting a nucleic acid-mediated interaction of RNA binding proteins with FAM60A.

### FAM60A-associated transcript codes for regulators of RNA processing

The observed attenuation of nucleic acid-mediated interactions post-nuclease treatment, as depicted in **Fig. 4D**, provides a compelling indication of FAM60A’s direct engagement with mRNA. To investigate this association, we first isolated FAM60A via affinity purification, then isolated the co-purified RNAs using RNA Affinity Purification (RAP) as elucidated in **Fig. 5A**. We overexpressed both Halo-FAM60A and FAM60A-Halo, along with Halo only, in HEK293 cells, followed by affinity purification of FAM60A (**Fig. 5B**). After purification, we extracted RNA and conducted sequencing on the mRNA co-purified with FAM60A (pulldown RNA) as well as on total the RNA samples taken from a portion of the cell lysate prior to protein isolation. This analysis revealed 368 mRNA entities significantly enriched in the FAM60A-affinity purified samples in comparison to control, satisfying the criteria of adjusted p-value < 0.05, a log2FC > 2, and mean FPKM>20 (**Fig. 5C** and **Suppl. Table S5**).

**Fig. 5:**
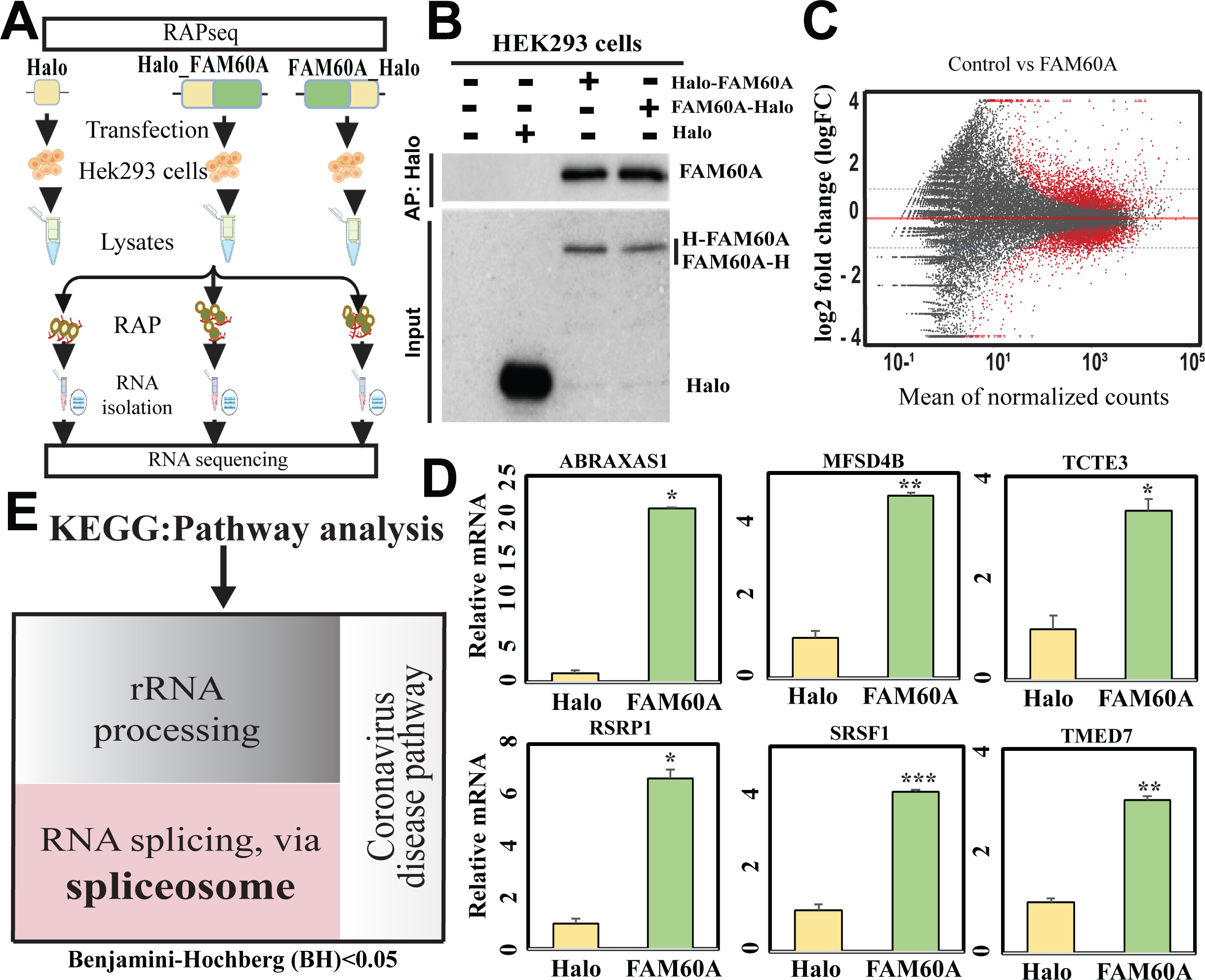
Sequencing analysis of mRNA copurifying with FAM60A. a, schematic-Diagram illustrating RNA affinity purification sequencing (RAPseq) for Halo-FAM60A and FAM60A-Halo. b, Halo-FAM60A or FAM60A-Halo was introduced into HEK293 cells, followed by affinity purification and immunoblot analysis to detect FAM60A presence. c, following RNA sequencing, mRNAs were identified as enriched in Halo-FAM60A and FAM60A-Halo and presented in MA plot (_adj_p < 0.05, log2 FC > 0.5, mean FPKM > 20). D, RT-qPCR was conducted to detect several mRNAs co-purify with FAM60A. e, pathway analysis of FAM60A-associated mRNAs showed enrichment in RNA processing and splicing, visualized using the tool RAVIGO. *P ≤ 0.05; **P ≤ 0.001; ***P ≤ 0.0001.

To corroborate the association of specific transcripts with FAM60A, we assessed six genes, including RSRP1 and SRSF1, known for their roles in RNA processing^25–27^, using qPCR. All evaluated genes showed results consistent with those observed in RAP-seq (**Fig. 5D**). Subsequent bioinformatics analysis of the 368 gene products enriched in the FAM60A-affinity purified samples using the DAVID functional annotation tool^28^ delineated 184 as protein-coding genes. Further analysis using the KEGG pathway analysis within DAVID elucidated significant enrichment in only three pathways, notably the coronavirus disease pathway, rRNA processing, and mRNA splicing via spliceosome pathways as visualized in RAVIGO^29^ (**Fig. 5E** and **Fig. S5A**). Our data strongly suggest that FAM60A is a pivotal player in the processing of RNA. Thus, these results underscore the necessity for further exploration into the consequences on RNA processing in the absence of FAM60A.

### FAM60A is a critical regulator of splicing has a global impact on cellular gene expression

We observed that FAM60A binds with both mRNA (**Fig. 5C**) and mRNA binding proteins (**Fig. 4A**), including spliceosome factors, suggesting its potential role in RNA splicing. Thus, to systematically evaluate this role, we employed CRISPR/Cas9 technology to generate a FAM60A knockout in HEK293 cell lines. The successful knockout was validated through RT-qPCR (**Fig. S6A**) and immunoblotting analyses (**Fig. 6A**). Comprehensive transcriptomic investigations of RNA from both knockout and control cell lines elucidated the crucial role of FAM60A in the regulation of RNA splicing, as delineated in **Fig. 6B** and **Suppl. Table S5**. We identified significant alterations in 139 splicing events demonstrating a notable change (≥ 15%) in FAM60A depleted cells, primarily in alternate first exon and exon skipping events (**Fig. 6B**). We observed additional splicing variations such as mutually exclusive exons, intron retention, and alterations at both alternative 5′ and 3′ splice sites (**Fig. 6B)**. To validate our findings, we used RT-PCR to confirm FAM60A was required for specific splice events, such as exon skipping, exon inclusion, intron retention, alternate first exon (**Fig. S6B**). For instance, in **Fig. 6C**, the inclusion of exon 23 of *SPTAN1* significantly increased in FAM60A-depleted cells, a result confirmed by RT-PCR and quantitative analysis. In contrast, in **Fig. 6D**, we observed a significant skipping of FN1 exon 25 in FAM60A-depleted cells compared to control, which was also validated by RT-PCR and quantitatively presented in a bar diagram (**Fig. 6D bottom**). Additionally, the impact of FAM60A knockout on alternate first exon splicing of *PPM1A* was confirmed through RT-PCR (**Fig. 6E**), while intron retention of *HNRNPA2B1* was also verified via RT-PCR and quantified, with results presented in a bar diagram (**Fig. S6C**).

**Fig. 6:**
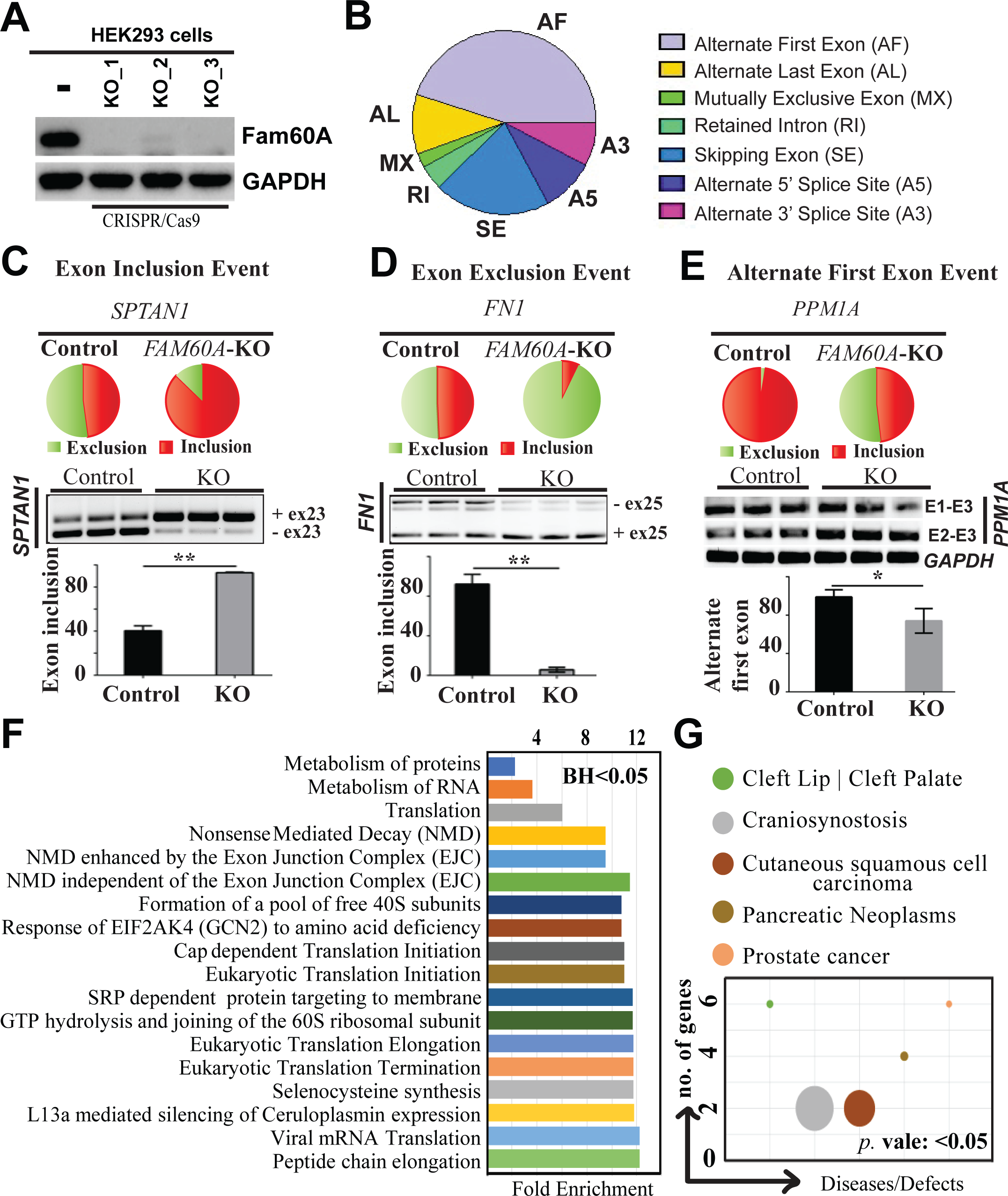
FAM60A involves in splicing. **a,** multiguide-CRISPR/cas9 was employed to Knocked out (KO) FAM60A in HEK 293 cells, lysates from FAM60A KO cells were analyzed by immunoblotting using antibodies against FAM60A, and GAPDH as a loading control. b, Pie chart illustrating 139 differentially regulated Alternate Splicing events (FDR≤ 5%, Δψ≥ 15%, n=3) between control and FAM60A knockout cells. c, d, and e, Pie charts displaying the Alternate Splicing events of exon inclusion, exclusion, and alternate first exon, which were validated and quantified through RT-PCR. F, bioinformatics analysis revealing that FAM60A impacts spliced mRNA, particularly in NMD pathways, among others. g, Demonstrating the disease associations related to the affected genes. *P ≤ 0.05; **P ≤ 0.001.

Pathway analysis executed through DAVID^28^ indicated that FAM60A plays a pivotal role in various pathways integral to RNA processing, metabolism, and mRNA degradation. Remarkably, FAM60A was identified as a critical factor for the splicing of pre-mRNAs, nonsense-mediated mRNA decay (NMD) pathway, encompassing both exon junction complex (EJC)-dependent and EJC-independent mechanisms (**Fig. 6F** and **Suppl. Table S5**). Concurrently, disease association studies conducted within the same analytical framework revealed a correlation between genes alternatively spliced via FAM60A and aberrant regulation in several diseases. These include cleft lip/palate, craniosynostosis, cutaneous squamous cell carcinoma, pancreatic neoplasms, and prostate cancer (**Fig. 6G** and **Suppl. Table S5**). This comprehensive analysis thus highlights FAM60A’s significant role in gene expression modulation and its potential implications in disease pathogenesis, offering novel insights for future therapeutic research endeavors.

## Discussion

FAM60A, an indispensable subunit of the Sin3/HDAC histone deacetylase complex, is essential in gene regulation and chromatin remodeling^6,7,9^. While its role within the Sin3/HDAC complex is well-established, FAM60A’s cellular functions beyond this specific complex remain largely uncharted. Notably, FAM60A plays a crucial role in maintaining stem cell pluripotency^7^, with its absence resulting in embryonic lethality^9^, indicating its broader, yet undefined, roles in cellular systems. Our comprehensive investigation into FAM60A’s molecular functional dynamics has uncovered: i) its integral association with the Sin3/HDAC complex; ii) FAM60A diverse structural domains that broaden its protein interactions beyond Sin3/HDAC; iii) its interaction with SIN3A, mediated through binding with SAP30 and HDAC1, as deciphered through advanced 3D modeling informed by cross-linking constraints; and iv) its role as an RNA-binding protein, significantly influencing RNA splicing processes. These insights collectively augment our comprehension of FAM60A’s central role in gene expression regulation, chromatin restructuring, and the sophisticated regulation of RNA splicing.

FAM60A is commonly recognized as a subunit of the Sin3/HDAC complex^6,7,14^, predominantly based on affinity-purification-MS (AP-MS) analyses that illustrate its interactions with Sin3/HDAC components. Our AP-MS investigations corroborate these findings, revealing reciprocal interactions between FAM60A and SIN3A, along with various other subunits (**Fig. 1B-D**). Notably, FAM60A utilizes its C-terminus to interact with the Sin3/HDAC complex and its N-terminus for associations with RNA and DNA binding proteins (**Fig. 2C-D**), demonstrating its dual functional capacity in molecular interactions. However, AP-MS analysis falls short in determining the direct binding partners of FAM60A within the Sin3/HDAC complex, particularly its interaction with SIN3A, the scaffold protein within the SIN3/HDAC complex^30,31^. To address this limitation, our research extends beyond proteomic analysis, using gel-filtration chromatography coupled with cross-linking proteomics. This approach elucidates the intricate framework of FAM60A’s direct interactions within the Sin3/HDAC complex. Contrary to expectations^6,7,9,14^, FAM60A does not bind directly with SIN3A. Instead, it forms the FAM60A-Sin3/HDAC complex through direct interactions with HDAC1 and SAP30 (**Fig. 3C-D**). Validation through CRISPR/Cas9-mediated knockouts of HDAC1 and SAP30, individually or simultaneously, confirmed that FAM60A’s primary association with the Sin3/HDAC complex is mediated through HDAC1 (**Fig. 3E**). This insight into FAM60A’s interaction dynamics enriches our understanding of its role in chromatin remodeling and gene regulation.

FAM60A plays a pivotal role in orchestrating the Sin3/HDAC complex’s genomic targeting and function within embryonic stem cells^7,9^. Our chemical cross-linking proteomics data unveil direct interactions between FAM60A and both HDAC1 and SAP30, which serve as bridges to SIN3A (**Fig. 3C and D**), refining the previously known role of FAM60A in complex assembly and genomic targeting. These findings enhance previous reports by demonstrating that FAM60A influences Sin3/HDAC complex binding to genomic regions through HDAC1 and SAP30 interactions. The disruption of FAM60A leads to embryonic lethality and promoter hypomethylation^9^, which we suggest arises from two distinct mechanisms. First, we discovered that FAM60A directly binds with MeCP2 (methyl-CpG-binding protein 2) (**Fig. 3C**), suggesting that FAM60A facilitates MeCP2 recruitment to methylated DNA sequences^32^ at gene promoters, thereby silencing gene expression. Without FAM60A, MeCP2’s binding to methylated DNA is hindered, making these regions susceptible to demethylation by transferases and leading to promoter hypomethylation. Second, FAM60A’s interaction with TET1, a demethylation enzyme^33^ observed specifically in pluripotent stem cells^7^, suggests another layer of spatiotemporal control. Collectively, our findings illuminate FAM60A as a master regulator of gene expression and chromatin remodeling, with its dysregulation leading to severe developmental defects.

Our comprehensive proteomic analysis significantly revises the existing perception of FAM60A, previously thought to be exclusively associated with the Sin3/HDAC complex^6^, revealing its ability to engage in a wider range of protein interactions. This extended functionality is facilitated by its distinct domains: the C-terminus facilitates interaction with the Sin3/HDAC complex (**Fig. 2C**), while the N-terminus, marked by a GATA-like zinc finger domain^8^, binds with both RNA and RNA binding proteins (**Fig. 4 A&D**), including spliceosome factors U2AF1, U2AF2, and SR proteins (SRSF1-SRSF7)^34–36^, particularly SRSF1^37^ and SRSF2^38^, which are involved in NMD^39^. Intriguingly, our study shows that FAM60A is actively involved in RNA splicing, and its depletion markedly impairs the splicing of pre-mRNAs linked to the NMD pathway, involving both EJC-dependent and EJC-independent mechanisms (**Fig. 6F**). These findings, especially the interaction of FAM60A with SRSF1 and SRSF2 associated with NMD, lay the groundwork for a future research program aimed at exploring FAM60A’s roles in mRNA decay and homeostasis, thereby enriching our understanding of its cellular functions.

In summary, our study redefines FAM60A’s cellular reach, extending its influence beyond the confines of the Sin3/HDAC complex. We reveal a versatile protein employing multiple domains to not only tether to the complex but also orchestrate a wider protein interactome. We dissect the mechanism by which FAM60A bridges the Sin3/HDAC complex through SAP30 and HDAC1, and unveil its independent role as an RNA-binding protein. Notably, FAM60A emerges as an important player in RNA processing, actively engaging with RNA molecules and influencing splicing events. This dual nature positions FAM60A as a critical regulator of gene expression, operating both at the chromatin and RNA levels. Our findings illuminate a previously uncharted landscape, inviting further exploration of FAM60A’s intricate interplay with RNA and its impact on cellular dynamics.

## Materials and Methods

### Materials

Magne® HaloTag® beads (G7281) and SNAP-Capture Magnetic Beads (S9145) were acquired from Promega (Madison, WI) and New England Biolabs (Ipswich, MA), respectively. HaloTag® TMRDirect™ Ligand (G2991) was sourced from Promega. AcTEV™ Protease (12575015) and PreScission Protease (27–0843-01) were obtained from Thermo Fisher Scientific and GE Health Life Sciences, respectively. Micrococcal Nuclease (M0247S) was procured from New England Biolabs. Antibodies used in this study were sourced as follows: RbAp46/RBBP7 (A22274), RBBP4 (A3645,), and SIN3A (A21966) from ABclonal, USA; Halo-Tag from Promega, USA (G9211); SAP30 (27679-1-AP) and SUDS3 (25845-1-AP) from Proteintech, USA; Fam60a from MRC PPU, Dundee University, UK (Cat # S381C). Rabbit anti-HaloTag® polyclonal antibody (G9281) was obtained from Promega. Mouse anti-tubulin monoclonal antibody (66031–1-Ig) came from ProteinTech. Goat anti-Mouse IRDye® 680LT (926–68020) and goat anti-Rabbit IRDye® 800CW (926–3211) secondary antibodies were obtained from LI-COR Biosciences. HRP-conjugated Goat Anti-Rabbit IgG(H+L) (SA00001-2), and Goat Anti-Mouse IgG(H+L) (SA00001-1) secondary antibodies were purchased from Proteintech, USA

### Cell culture

HEK293 and 293T cells were cultured in Dulbecco’s Modified Eagle’s Medium (DMEM), enriched with 10% fetal bovine serum (FBS) and 2 mM Glutamax. The cells were incubated at a temperature of 37°C in an incubator with 5% CO_2_.

### Construction of Expression Vectors

Expression vectors were generated following previously described methodologies^15,40^. Briefly, target genes were subcloned into the Halo pcDNA5/FRT vector, specifically between the PacI and PmeI restriction sites. This vector construction was subsequently verified through sequencing. For the production of Halo-tagged constructs expressing Fam60A and SIN3A deletion mutants, the corresponding open reading frames (ORFs) were PCR-amplified and subcloned into the Halo pcDNA5/FRT vector using Gibson Assembly Cloning Kit [New England Biotechnologies (NEB)]. Specifically, for the creation of a C-terminal Halo-tagged construct of Fam60A (Fam60A-Halo) in the pcDNA5/FRT system, the ORF of Fam60A was amplified using PCR and then subcloned into the pcDNA5FRT C-Halo construct. All plasmid constructs were confirmed by sequencing. The primers used for cloning and PCR are listed in **Suppl. Table S6**.

### Preparation of cell lysates

Cells that were confluent or nearly confluent were collected and rinsed twice with ice-cold PBS. Unless otherwise stated, the entire processes were performed at 4°C (or in an iced environment). Cells were then suspended in a freshly made lysis buffer (containing 20 mM Tris with pH 7.5, 1% triton, 150 mm NaCl, and protease inhibitors: Aprotinin at 5 mg/l and PMSF at 0.1 mM) and left on ice for a half-hour. Then, the cells were lysed by using 26 G syringe by repeating several times. This was followed by centrifuging at 14,000 rpm for 15 minutes at 4°C.

### Live cell imaging

For this experiment, HEK 293 cells were planted at 20% confluency on glass-bottomed culture dishes (supplied by MatTek, Ashland, MA: (35mm, No. 2 14-mm diameter glass) and then transiently transfected with Halo-tagged Fam60A/SIN3A and Halo-tagged mutants Fam60A/SIN3A constructs. During growth, affinity-tagged proteins were fluorescently marked, with Halo-Tag TMRDirect ligand (Promega, WI, USA), as per the manufacturer’s guidelines. Images were captured using a Zeiss LSM 780 confocal microscope, with argon laser excitation at 573-687nm for TMRDirect. To prevent photobleaching, exposure duration and laser power were tweaked to improve image quality. An alternating excitation mode was implemented to prevent cross-talk among color channels. The HaloTag™ TMRDirect ligand was added to label HaloTag™ proteins at a final concentration of 100 nM, and the cells were incubated overnight at 37°C in 5% CO_2_. The culture medium was swapped for OptiMEM to minimize background fluorescence before imaging. Cells were stained with Hoeschst dye for 30 minutes to identify nuclei before imaging.

### Halo affinity purification for proteomic analysis

HEK293T cells (1 × 10^7^) were cultured in 15-cm plates for 24 hours. They were then transfected with Halo-tagged gene constructs using Lipofectamine LTX (Thermo Fisher Scientific, MA, USA). After 48 hours, cells were harvested and washed twice with ice-cold PBS. The cells were lysed in 300 μl of mammalian cell lysis buffer (Promega, WI, USA), containing 50 mM Tris-HCl (pH 7.5), 150 mM NaCl, 1% Triton X-100, 0.1% sodium deoxycholate, 0.1 mM benzamidine HCl, 55 μM phenanthroline, 1 mM PMSF, 10 μM bestatin, 5 μM pepstatin A, and 20 μM leupeptin. Lysed cells were passed through a 26-gauge needle 5-7 times and then centrifuged at 14,000rpm for 30 minutes at 4°C. The supernatant (300 μl) was mixed with 700 μl of Tris-buffered saline and centrifuged again at 14,000rpm for 10 minutes at 4°C. The clear supernatant was incubated overnight at 4°C with 100 μl of prewashed Magne® HaloTag® Beads (Promega, WI, USA). Post-incubation, the beads were washed four times with 750 μl of wash buffer and eluted with a buffer containing 50 mM Tris-HCl (pH 8.0), 0.5 mM EDTA, 0.005 mM DTT, and 2 Units of AcTEV Protease for 2 hours at room temperature. Finally, the eluate was passed through a Micro Bio-Spin column to remove any bead residues before proteomic analysis.

### MudPIT analysis for proteins and protein complexes identification

The procedure for identifying protein complexes using MudPIT analysis has been previously detailed by Banks et al.,^40^. To provide a concise summary, proteins purified through trichloroacetic acid precipitation were first subjected to proteolytic digestion with endoproteinase Lys-C, followed by trypsin digestion. This process was carried out overnight at a temperature of 37°C. Subsequently, a 10-step MudPIT separation technique was utilized, and the digested peptides were directly injected into a linear ion trap mass spectrometer for the collection and identification of spectra. The analysis of peptide mass spectra was performed using the ProLuCID and DTASelect algorithms. We then implemented Contrast and NSAF7 software to arrange the potential affinity-purified proteins based on their respective distributed normalized spectral abundance values. To identify proteins in the experimental samples that were more abundant compared to the control samples, we utilized the QSPEC software. False discovery rates from QSPEC parameters appropriate for multiple comparisons were computed using the Benjamini-Hochberg statistical method. Unless specified otherwise, each experiment was conducted at least three times. All raw mass spectrometry data are accessible as described in **Suppl. Table S6**.

### Immunoblotting

Protein lysates were separated on 10% SDS-PAGE gels and then transferred to PVDF membranes (GE Healthcare Life Science). The membranes were blocked in phosphate-buffered saline with Tween 20 (PBST) containing 5% BSA for 1 h at room temperature. Thereafter, membranes were incubated with respective primary antibodies overnight at 4 °C. After washing three times with Tris Buffered Saline with Tween 20 (TBST) for 5 min each, the membranes were incubated with horseradish peroxidase-conjugated secondary antibodies at room temperature for 2 h. The membranes were washed three times with TBST. The specific proteins were visualized using ECL western blotting system (ThermoFisher, USA). For quantitation of western blot band intensity, densitometric analysis was performed using Image J software and normalized to the loading control. For SIN3A and mutants experiments, membranes were probed with respective primary antibodies, followed by incubation with IRDye® 680LT Goat-anti-Mouse, IRDye® 800CW Goat-anti-Mouse, or IRDye® 800CW Goat-anti-Rabbit secondary antibodies (LI-COR), all diluted 1:10,000. Images of the blots were captured using an Odyssey® CLx imaging system (LI-COR).

### Gel Filtration chromatography

Purified Fam60A protein complexes were isolated from whole cell extracts of HEK293T cells expressing Halo-Fam60A using the Halo affinity purification method. The purified Fam60A was then subjected to size-exclusion chromatography on a Superose 6, 10/300 GL column (Amersham Bioscience) using a buffer composed of 40mM HEPES (pH 7.9), 350mM NaCl, 5% glycerol, 0.1% Tween 20, and 1.0mM DTT. Fractions (500μl each) were collected to analyze the co-fractionation profiles of Fam60A and its associated proteins, identified through LC-MS/MS (MudPIT). The column calibration was performed using a set of gel filtration markers (Sigma-Aldrich cat. # MWGF1000), including Blue Dextran 2000, thyroglobulin (669kDa), Ferritin (440kDa), β-amylase (200kDa), Alcohol dehydrogenase (150), and bovine serum albumin (66kDa).

### XL-MS analysis

The preparation of samples for cross-link proteomics was outlined previously^15,19^. Briefly, cell lysates expressing Halo-Fam60A were mixed with Magne® HaloTag® Beads, using 100 μl of bead slurry, and incubated overnight at 4°C, following the manufacturer’s protocol. The beads were then separated with a magnetic concentrator and washed four times using a buffer composed of 10 mM HEPES (pH 7.5), 1.5 mM MgCl2, 0.3 M NaCl, 10 mM KCl, and 0.2% Triton X-100. To elute bound proteins, the beads were treated with 200 μl of a buffer containing 50 mM HEPES (pH 7.5), 0.5 mM EDTA, 1 mM DTT, and 30 units of AcTEV, and incubated for a minimum of 2 hours at 4°C. The eluate was collected, and proteins were cross-linked using 5 mM DSSO for 40 minutes at room temperature. This cross-linking process was stopped by adding 50 mM NH_4_HCO_3_ and incubating for 15 minutes at room temperature.

### RNA Pull-down Assay

As we described previously^41^, in RNA pull-down assay, approximately 100 million transiently transfected HEK293T cells were lysed in 300 μl of ice-cold buffer containing, 50 mm Tris·HCl, 150 mm NaCl, and 1% Triton® X-100. The lysate was passed through a 26-gauge needle five times, centrifuged at 14,000 rpm at 4 °C for 30 minutes, and the supernatant was diluted with 700 μl TBS. Protein complexes were isolated from 900 μl of the cell extract using Magne® HaloTag® Beads and eluted with AcTEV™ Protease and RNaseOUT™. RNA was extracted from these samples or from the full cell extract using RNeasy® Mini kits, with DNA removal using DNase I. The RNA was reverse transcribed and sequenced, aligning with the UCSC hg19 genome.

### Knockout cell generation

For the knockout of the *FAM60A* gene using the CRISPR/Cas9 system, multiple high-specificity guide RNAs (gRNAs) targeting three distinct regions of Fam60A were designed, synthesized, and inserted into a CRISPR/Cas9 vector^42^. This construct was then transfected into the HEK293 cell line. Following transfection, cells underwent puromycin selection were selected to establish a homogenous pool of cells with disrupted Fam60A. The efficiency of Fam60A knockout was confirmed through Western blot and qRT-PCR analyses.

### qRT-PCR and RT-PCR

Following the instructions of the manufacturer, total RNA was extracted using the Quick-RNA™ Miniprep Kit, incorporating DNase-I treatment to eliminate host DNA contamination during extraction process. For gene expression assessment, a two-step qRT-PCR approach was used which targeted both the gene of interest and a housekeeping gene (GAPDH) using SsoAdvanced™ Universal SYBR® Green Supermix (BIO-RAD, CA, USA). First, reverse transcription of 1µg RNA was executed with the iScript™ cDNA Synthesis Kit (BIO-RAD, CA, USA). Then quantitative PCR reaction was performed in 20μl final volumes, consisted of 1 × SsoAdvanced™ Universal SYBR® Green Supermix, 500nM each of forward and reverse primers, 10 ng cDNA, and nuclease-free water. The CFX Opus 96 Real-Time PCR System (Bio-Rad Laboratories, CA, USA) was utilized to determine the relative mRNA expression levels. In the RT-PCR analysis, cDNA was amplified using specific primers for the target genes. The resulting PCR products were then separated on an agarose gel for visualization and confirmation of amplification. Band intensities were subsequently quantified to analyze gene expression with a minimum of three repetitions for all PCR experiments.

### Detecting and Validating alternative splicing events

Genome-wide analysis of differential alternative splicing events was conducted using the SUPPA2 tool^43^. To delineate significant splicing variations, we employed criteria such as the inclusion and exclusion read counts exhibiting a |Δψ| ≥ 15%, along with an FDR-adjusted p-value < 0.05. For qRT-PCR, primers targeting an intronic and exonic region, as well as primers spanning various splicing junctions, were utilized for mRNAs from both control and Fam60A knockout (KO) cells (**Suppl. Table S6**).

### Statistical analysis

For the analysis of quantitative PCR (qPCR) data involving multiple comparisons, we employed a student’s t-test or one-way analysis of variance (ANOVA) followed by a Newman-Keuls post hoc test where appropriate. This statistical analysis was conducted using GraphPad Prism software, version 5.04 (GraphPad Software, San Diego, CA, USA). We presented the results as means ± standard deviation (SD), with a sample size of n ≥ 3, unless specified otherwise. A p-value of 0.05 or less (P ≤ 0.05) was considered to indicate statistical significance.

## Acknowledgements

Our sincere thanks go to Charles A. S. Banks for his invaluable assistance in preparing figures and generating plasmids, and to Mihaela Sardiu for insightful discussions on data analysis strategies for this study.

## Author contributions

S.M. conceived the study. S.M., M.N.H., C.A.S.B., M.A.R., L.F., and M.P.W. designed experiments and interpreted all experiments. S.M., M.N.H., C.A.S.B., R.Z., P.N., J.A., M.R.I., M.M., C.G.K., J.L.T., Y.Z., Z.W., G.H., A. G., and G.D. performed experiments. S.M., M.N.H., C.A.S.B., R.Z., P.N., M.A.R., S.B., L.F., and M.P.W. analyzed data and interpreted findings. S.M. R.Z., M.N.H., and M.M. wrote the first draft and L.F., M.A.R., and M.P.W. revised the manuscript. S.M., L.F., and M.P.W. guided the research.

## Data and materials availability

All data supporting the paper’s conclusions are available in the paper and its Supplementary Materials. Materials inquiry requests should be made to mwashburn4{at}kumc.edu and msmiah{at}uams.edu.

## References

1. Kouzarides, T. (2007). Chromatin Modifications and Their Function. Cell 128, 693–705. 10.1016/j.cell.2007.02.005.

2. Alam, J., Huda, M.N., Tackett, A.J., and Miah, S. (2023). Oncogenic signaling-mediated regulation of chromatin during tumorigenesis. Cancer Metastasis Rev. 42, 409–425. 10.1007/s10555-023-10104-3.

3. Lai, A.Y., and Wade, P.A. (2011). Cancer biology and NuRD: A multifaceted chromatin remodelling complex. Nat. Rev. Cancer 11, 588–596. 10.1038/nrc3091.

4. Arnould, C., Rocher, V., Saur, F., Bader, A.S., Muzzopappa, F., Collins, S., Lesage, E., Le Bozec, B., Puget, N., Clouaire, T., et al. (2023). Chromatin compartmentalization regulates the response to DNA damage. Nature 623, 183–192. 10.1038/s41586-023-06635-y.

5. Kadamb, R., Mittal, S., Bansal, N., Batra, H., and Saluja, D. (2013). Sin3 : Insight into its transcription regulatory functions. Eur. J. Cell Biol. 92, 237–246. 10.1016/j.ejcb.2013.09.001.

6. Smith, K.T., Sardiu, M.E., Martin-Brown, S.A., Seidel, C., Mushegian, A., Egidy, R., Florens, L., Washburn, M.P., and Workman, J.L. (2012). Human family with sequence similarity 60 member A (FAM60A) protein: A new subunit of the Sin3 deacetylase complex. Mol. Cell. Proteomics 11, 1815–1828. 10.1074/mcp.M112.020255.

7. Streubel, G., Fitzpatrick, D.J., Oliviero, G., Scelfo, A., Moran, B., Das, S., Munawar, N., Watson, A., Wynne, K., Negri, G.L., et al. (2017). Fam60a defines a variant Sin3a-Hdac complex in embryonic stem cells required for self-renewal. EMBO J. 10.15252/embj.201696307.

8. Muñoz, I.M., MacArtney, T., Sanchez-Pulido, L., Ponting, C.P., Rocha, S., and Rouse, J. (2012). Family with sequence similarity 60A (FAM60A) protein is a cell cycle-fluctuating regulator of the SIN3-HDAC1 histone deacetylase complex. J. Biol. Chem. 287, 32346–32353. 10.1074/jbc.M112.382499.

9. Nabeshima, R., Nishimura, O., Maeda, T., Shimizu, N., Ide, T., Yashiro, K., Sakai, Y., Meno, C., Kadota, M., Shiratori, H., et al. (2018). Loss of Fam60a, a Sin3a subunit, results in embryonic lethality and is associated with aberrant methylation at a subset of gene promoters. Elife 7, 1–29. 10.7554/eLife.36435.

10. Hou, Q., Jiang, Z., Li, Y., Wu, H., Yu, J., and Jiang, M. (2020). FAM60A promotes cisplatin resistance in lung cancer cells by activating SKP2 expression. Anticancer. Drugs, 776–784. 10.1097/CAD.0000000000000952.

11. Yue, X., Xi, X., Cheng, B., and Chen, Y. (2023). SIN3A/HDAC complex subunit FAM60A is associated with proliferation in colorectal cancer. Asian J. Surg., 1015–1016. 10.1016/j.asjsur.2023.01.045.

12. Yan, Z., Jun-Hui, L., Yuan, Q.-G., and Yang, W.-B. (2022). Restraint of FAM60A has a cancer-inhibiting role in pancreatic carcinoma via the effects on the Akt/GSK-3β/β-catenin signaling pathway. Environ. Toxicol. 37, 1432–1444. 10.1002/tox.23496.

13. Smith, K.T., Sardiu, M.E., Martin-Brown, S. a., Seidel, C., Mushegian, a., Egidy, R., Florens, L., Washburn, M.P., and Workman, J.L. (2012). Functional characterization of the human FAM60A protein: a new subunit of the Sin3 deacetylase complex. Mol. Cell. Proteomics, 1815–1828. 10.1074/mcp.M112.020255.

14. Sardiu, M.E., Smith, K.T., Groppe, B.D., Gilmore, J.M., Saraf, a., Egidy, R., Peak, a., Seidel, C.W., Florens, L., Workman, J.L., et al. (2014). Suberoylanilide Hydroxamic Acid (SAHA)-Induced Dynamics of a Human Histone Deacetylase Protein Interaction Network. Mol. Cell. Proteomics 13, 3114–3125. 10.1074/mcp.M113.037127.

15. Adams, M.K., Banks, C.A.S., Thornton, J.L., Kempf, C.G., Zhang, Y., Miah, S., Hao, Y., Sardiu, M.E., Killer, M., Hattem, G.L., et al. (2020). Differential Complex Formation via Paralogs in the Human Sin3 Protein Interaction Network. Mol. Cell. Proteomics 19, 1468–1484. 10.1074/mcp.RA120.002078.

16. Shannon, P., Markiel, A., Ozier, O., Baliga, N.S., Wang, J.T., Ramage, D., Amin, N., Schwikowski, B., and Ideker, T. (2003). Cytoscape: A Software Environment for Integrated Models of biomolecular interaction networks. Genome Res. 13, 426. doi: 10.1101/gr.1239303.

17. Sturn, A., Quackenbush, J., and Trajanoski, Z. (2002). Genesis: cluster analysis of microarray data. Bioinformatics 18, 207–208. 10.1093/bioinformatics/18.1.207.

18. Sturn, A., Mlecnik, B., Pieler, R., Rainer, J., Truskaller, T., and Trajanoski, Z. (2003). Client-server environment for high-performance gene expression data analysis. Bioinformatics 19, 772– 773. 10.1093/bioinformatics/btg074.

19. Banks, C.A.S., Zhang, Y., Miah, S., Hao, Y., Adams, M.K., Wen, Z., Thornton, J.L., Florens, L., and Washburn, M.P. (2020). Integrative Modeling of a Sin3/HDAC Complex Sub-structure. Cell Rep. 31. 10.1016/j.celrep.2020.03.080.

20. Jumper, J., Evans, R., Pritzel, A., Green, T., Figurnov, M., Ronneberger, O., Tunyasuvunakool, K., Bates, R., Žídek, A., Potapenko, A., et al. (2021). Highly accurate protein structure prediction with AlphaFold. Nature 596, 583–589. 10.1038/s41586-021-03819-2.

21. Zhou, X., Zheng, W., Li, Y., Pearce, R., Zhang, C., Bell, E.W., Zhang, G., and Zhang, Y. (2022). I-TASSER-MTD: a deep-learning-based platform for multi-domain protein structure and function prediction. Nat. Protoc. 17, 2326–2353. 10.1038/s41596-022-00728-0.

22. Merkley, E.D., Rysavy, S., Kahraman, A., Hafen, R.P., Daggett, V., and Adkins, J.N. (2014). Distance restraints from crosslinking mass spectrometry: Mining a molecular dynamics simulation database to evaluate lysine-lysine distances. Protein Sci. 23, 747–759. 10.1002/pro.2458.

23. Abdelmohsen, K., and Gorospe, M. (2012). RNA-binding protein nucleolin in disease. RNA Biol. 9, 799–808. 10.4161/rna.19718.

24. Doron-Mandel, E., Koppel, I., Abraham, O., Rishal, I., Smith, T.P., Buchanan, C.N., Sahoo, P.K., Kadlec, J., Oses-Prieto, J.A., Kawaguchi, R., et al. (2021). The glycine arginine-rich domain of the RNA-binding protein nucleolin regulates its subcellular localization. EMBO J. 40, 1–22. 10.15252/embj.2020107158.

25. Li, Y., Wang, X., Qi, S., Gao, L., Huang, G., Ren, Z., Li, K., Peng, Y., Yi, G., Guo, J., et al. (2021). Spliceosome-regulated RSRP1-dependent NF-κB activation promotes the glioblastoma mesenchymal phenotype. Neuro. Oncol. 23, 1693–1708. 10.1093/neuonc/noab126.

26. Saltzman, A.L., Pan, Q., and Blencowe, B.J. (2011). Regulation of alternative splicing by the core spliceosomal machinery. Genes Dev. 25, 373–384. 10.1101/gad.2004811.

27. Anczuków, O., Akerman, M., Cléry, A., Wu, J., Shen, C., Shirole, N.H., Raimer, A., Sun, S., Jensen, M.A., Hua, Y., et al. (2015). SRSF1-Regulated Alternative Splicing in Breast Cancer. Mol. Cell 60, 105–117. 10.1016/j.molcel.2015.09.005.

28. Sherman, B.T., Hao, M., Qiu, J., Jiao, X., Baseler, M.W., Lane, H.C., Imamichi, T., and Chang, W. (2022). DAVID: a web server for functional enrichment analysis and functional annotation of gene lists (2021 update). Nucleic Acids Res. 50, W216–W221. 10.1093/nar/gkac194.

29. Supek, F., Bošnjak, M., Škunca, N., and Šmuc, T. (2011). Revigo summarizes and visualizes long lists of gene ontology terms. PLoS One 6. 10.1371/journal.pone.0021800.

30. Grzenda, A., Lomberk, G., Zhang, J.-S., and Urrutia, R. (2009). Sin3 Master Scaffold and Transcriptional Corepressor. Biochim Biophys Acta. 0, 443–450. 10.1016/j.bbagrm.2009.05.007.

31. Pantier, R., Mullin, N.P., and Chambers, I. (2017). A new twist to Sin3 complexes in pluripotent cells. EMBO J. 36, 2184–2186. 10.15252/embj.201797516.

32. Jones, P.L., Veenstra, G.J.C., Wade, P.A., Vermaak, D., Kass, S.U., Landsberger, N., Strouboulis, J., and Wolffe, A.P. (1998). Methylated DNA and MeCP2 recruit histone deacetylase to repress transcription. Nat. Genet. 19, 187–191. 10.1038/561.

33. Liu, W., Wu, G., Xiong, F., and Chen, Y. (2021). Advances in the DNA methylation hydroxylase TET1. Biomark. Res. 9, 1–12. 10.1186/s40364-021-00331-7.

34. Rahman, M.A., Azuma, Y., Nasrin, F., Takeda, J.I., Nazim, M., Ahsan, K. Bin, Masuda, A., Engel, A.G., and Ohno, K. (2015). SRSF1 and hnRNP H antagonistically regulate splicing of COLQ exon 16 in a congenital myasthenic syndrome. Sci. Rep. 5, 1–13. 10.1038/srep13208.

35. Rahman, M.A., Krainer, A.R., and Abdel-Wahab, O. (2020). SnapShot: Splicing Alterations in Cancer. Cell 180, 208–208.e1. 10.1016/j.cell.2019.12.011.

36. Long, J.C., and Caceres, J.F. (2009). The SR protein family of splicing factors: Master regulators of gene expression. Biochem. J. 417, 15–27. 10.1042/BJ20081501.

37. Aznarez, I., Nomakuchi, T.T., Tetenbaum-Novatt, J., Rahman, M.A., Fregoso, O., Rees, H., and Krainer, A.R. (2018). Mechanism of Nonsense-Mediated mRNA Decay Stimulation by Splicing Factor SRSF1. Cell Rep. 23, 2186–2198. 10.1016/j.celrep.2018.04.039.

38. Rahman, M.A., Lin, K.T., Bradley, R.K., Abdel-Wahab, O., and Krainer, A.R. (2020). Recurrent SRSF2 mutations in MDS affect both splicing and NMD. Genes Dev. 34, 413–427. 10.1101/gad.332270.119.

39. Maquat, L.E. (2004). NONSENSE-MEDIATED mRNA DECAY : SPLICING, TRANSLATION AND mRNP DYNAMICS. Nature 5. 10.1038/nrm1310.

40. Banks, C.A.S., Thornton, J.L., Eubanks, C.G., Adams, M.K., Miah, S., Boanca, G., Liu, X., Katt, M.L., Parmely, T.J., Florens, L., et al. (2018). A structured workflow for mapping human sin3 histone deacetylase complex interactions using halo-MudPIT affinity-purification mass spectrometry. Mol. Cell. Proteomics 17, 1432–1447. 10.1074/mcp.TIR118.000661.

41. Banks, C.A.S., Boanca, G., Lee, Z.T., Eubanks, C.G., Hattem, G.L., Peak, A., Weems, L.E., Conkright, J.J., Florens, L., and Washburn, M.P. (2016). TNIP2 is a Hub Protein in the NF-κB Network with Both Protein and RNA Mediated Interactions. Mol. Cell. Proteomics 15, 3435– 3449. 10.1074/mcp.M116.060509.

42. Cao, J., Wu, L., Zhang, S.M., Lu, M., Cheung, W.K.C., Cai, W., Gale, M., Xu, Q., and Yan, Q. (2016). An easy and efficient inducible CRISPR/Cas9 platform with improved specificity for multiple gene targeting. Nucleic Acids Res. 44, 1–10. 10.1093/nar/gkw660.

43. Trincado, J.L., Entizne, J.C., Hysenaj, G., Singh, B., Skalic, M., Elliott, D.J., and Eyras, E. (2018). SUPPA2: Fast, accurate, and uncertainty-aware differential splicing analysis across multiple conditions. Genome Biol. 19, 1–11. 10.1186/s13059-018-1417-1.

