## Supplemental Figures for "Beyond the Sin3/HDAC Complex: FAM60A emerges as a regulator of RNA Splicing"

### Fig. S1

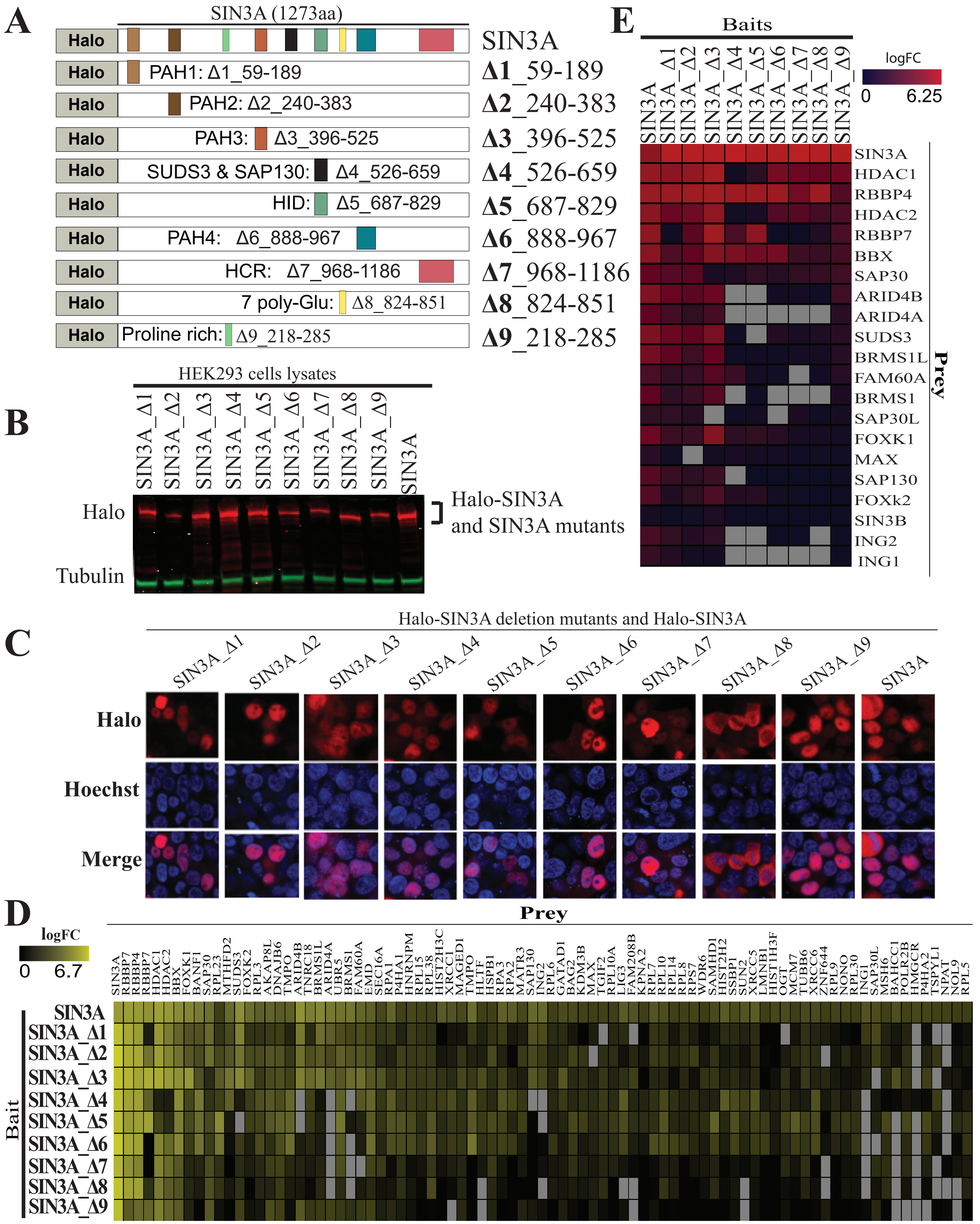
A
SIN3A (1273aa)
Halo
SIN3A
PAH1: Δ1_59-189
Δ1_59-189
PAH2: Δ2_240-383
Δ2_240-383
PAH3: Δ3_396-525
Δ3_396-525
SUDS3 & SAP130: Δ4_526-659
Δ4_526-659
HID: Δ5_687-829
Δ5_687-829
PAH4: Δ6_888-967
Δ6_888-967
HCR: Δ7_968-1186
Δ7_968-1186
7 poly-Glu: Δ8_824-851
Δ8_824-851
Proline rich: Δ9_218-285
Δ9_218-285
B
HEK293 cells lysates
SIN3A_Δ1 SIN3A_Δ2 SIN3A_Δ3 SIN3A_Δ4 SIN3A_Δ5 SIN3A_Δ6 SIN3A_Δ7 SIN3A_Δ8 SIN3A_Δ9 SIN3A
Halo
Tubulin
Halo-SIN3A
and SIN3A mutants
E
Baits
SIN3A SIN3A_Δ1 SIN3A_Δ2 SIN3A_Δ3 SIN3A_Δ4 SIN3A_Δ5 SIN3A_Δ6 SIN3A_Δ7 SIN3A_Δ8 SIN3A_Δ9
logFC
0 6.25
SIN3A HDAC1 RBBP4 HDAC2 RBBP7 BBX SAP30 ARID4B ARID4A SUDS3 BRMS1L FAM60A BRMS1 SAP30L FOXK1 MAX SAP130 FOXk2 SIN3B ING2 ING1
Prey
C
Halo-SIN3A deletion mutants and Halo-SIN3A
SIN3A_Δ1 SIN3A_Δ2 SIN3A_Δ3 SIN3A_Δ4 SIN3A_Δ5 SIN3A_Δ6 SIN3A_Δ7 SIN3A_Δ8 SIN3A_Δ9 SIN3A
Halo
Hoechst
Merge
D
logFC
0 6.7
Prey
SIN3A RBBP7 RBBP4 RBBP7 HDAC1 HDAC2 BBX FOXK1 BANF1 SAP30 RPL23 MTHFD2 SUDS3 FOXK2 RPL3 AKAP8L DNAJB6 TMPO ARID4B TNRC18 BRMS1L ARID4A UBR5 BRMS1 FAM60A EMD SEC16A RPA1 P4HA1 HNRNPM RPL15 RPL38 HIST2H3C XRCC1 MAGED1 TMPO HLTF HSPB1 RPA3 RPA2 MATR3 SAP130 ING2 RPL6 GATAD1 BAG2 KDM3B MAX TGIF2 RPL10A LIG3 FAM208B KPNA2 RPL7 RPL10 RPL14 RPL8 RPS7 WDR6 SAMHD1 HIST2H2 SSBP1 SUN2 XRCC5 LMNB1 HIST1H3F OGT MCM7 TUBB6 XRCC6 ZNF644 RPL9 NONO RPL30 ING1 SAP30L MSH6 BAHCC1 POLR2B HMGCR P4HA2 TSPYL1 NPAT NOL9 RPL5
Bait
SIN3A SIN3A_Δ1 SIN3A_Δ2 SIN3A_Δ3 SIN3A_Δ4 SIN3A_Δ5 SIN3A_Δ6 SIN3A_Δ7 SIN3A_Δ8 SIN3A_Δ9

Fig. S2

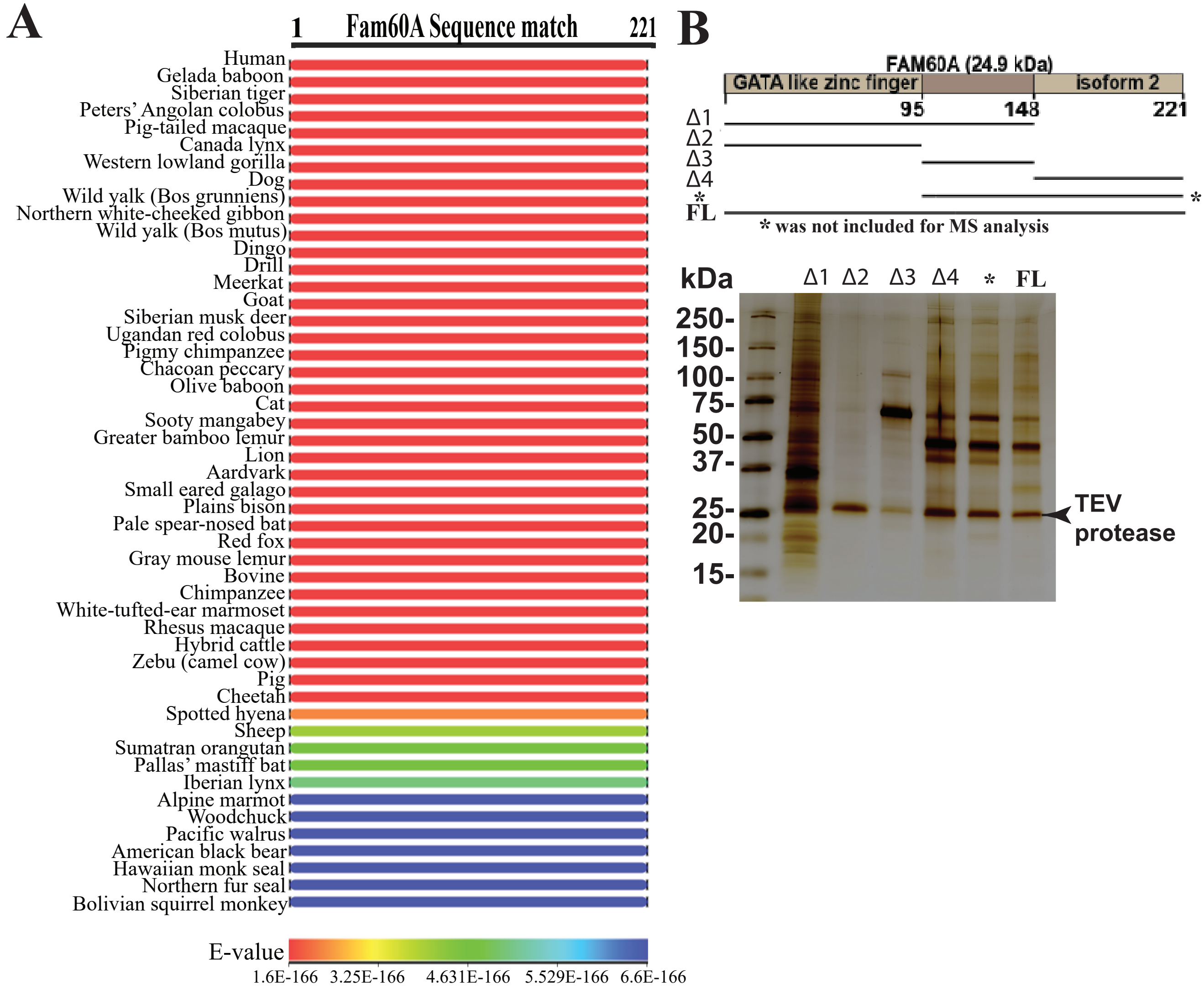
A
1
Fam60A Sequence match
221
Human
Gelada baboon
Siberian tiger
Peters' Angolan colobus
Pig-tailed macaque
Canada lynx
Western lowland gorilla
Dog
Wild yalk (Bos grunniens)
Northern white-cheeked gibbon
Wild yalk (Bos mutus)
Dingo
Drill
Meerkat
Goat
Siberian musk deer
Ugandan red colobus
Pigmy chimpanzee
Chacoan peccary
Olive baboon
Cat
Sooty mangabey
Greater bamboo lemur
Lion
Aardvark
Small eared galago
Plains bison
Pale spear-nosed bat
Red fox
Gray mouse lemur
Bovine
Chimpanzee
White-tufted-ear marmoset
Rhesus macaque
Hybrid cattle
Zebu (camel cow)
Pig
Cheetah
Spotted hyena
Sheep
Sumatran orangutan
Pallas' mastiff bat
Iberian lynx
Alpine marmot
Woodchuck
Pacific walrus
American black bear
Hawaiian monk seal
Northern fur seal
Bolivian squirrel monkey
E-value
1.6E-166
3.25E-166
4.631E-166
5.529E-166
6.6E-166
B
FAM60A (24.9 kDa)
GATA like zinc finger
isoform 2
1
95
148
221
Δ1
Δ2
Δ3
Δ4
*
FL
*
* was not included for MS analysis
kDa
Δ1
Δ2
Δ3
Δ4
*
FL
250-
150-
100-
75-
50-
37-
25-
20-
15-
TEV
protease

Fig. S3

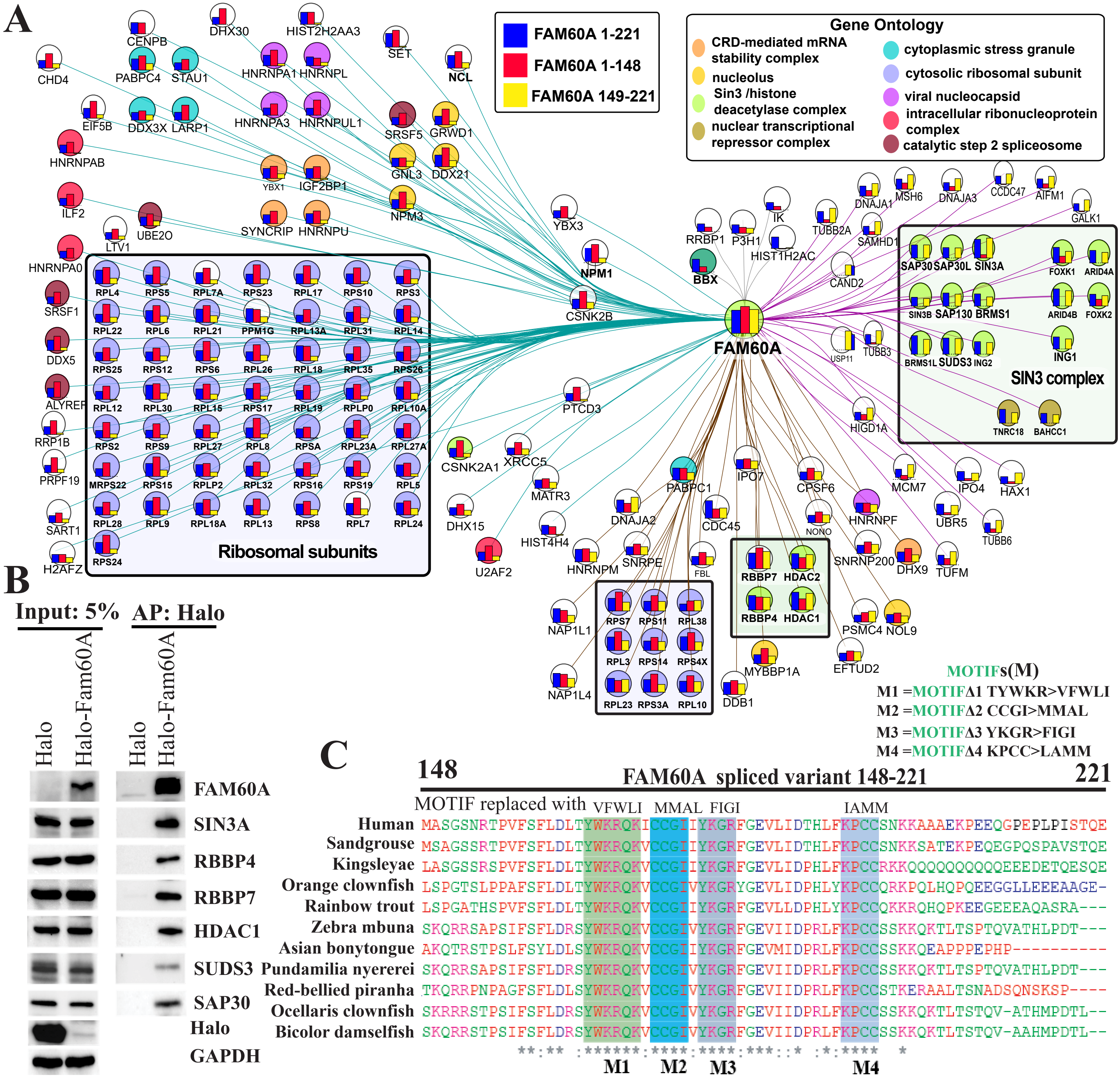
A
FAM60A 1-221
FAM60A 1-148
FAM60A 149-221
Gene Ontology
CRD-mediated mRNA stability complex
nucleolus
Sin3 /histone deacetylase complex
nuclear transcriptional repressor complex
cytoplasmic stress granule
cytosolic ribosomal subunit
viral nucleocapsid
intracellular ribonucleoprotein complex
catalytic step 2 spliceosome
FAM60A
Ribosomal subunits
SIN3 complex
CHD4
CENPB
DHX30
HIST2H2AA3
SET
NCL
PABPC4
STAU1
HNRNPA1
HNRNPL
EIF5B
DDX3X
LARP1
HNRNPA3
HNRNPUL1
SRSF5
GRWD1
HNRNPAB
YBX1
IGF2BP1
GNL3
DDX21
ILF2
UBE2O
SYNCRIP
HNRNPU
NPM3
YBX3
LTV1
HNRNPA0
NPM1
CSNK2B
SRSF1
DDX5
ALYREF
RRP1B
PRPF19
SART1
H2AFZ
RPL4
RPS5
RPL7A
RPS23
RPL17
RPS10
RPS3
RPL22
RPL6
RPL21
PPM1G
RPL13A
RPL31
RPL14
RPS25
RPS12
RPS6
RPL26
RPL18
RPL35
RPS26
RPL12
RPL30
RPL15
RPS17
RPL19
RPLP0
RPL10A
RPS2
RPS9
RPL27
RPL8
RPSA
RPL23A
RPL27A
MRPS22
RPS15
RPLP2
RPL32
RPS16
RPS19
RPL5
RPL28
RPL9
RPL18A
RPL13
RPS8
RPL7
RPL24
RPS24
PTCD3
CSNK2A1
XRCC5
MATR3
DHX15
HIST4H4
U2AF2
HNRNPM
SNRPE
DNAJA2
PABPC1
IPO7
CDC45
FBL
CPSF6
NONO
HNRNPF
SNRNP200
DHX9
TUFM
RBBP7
HDAC2
RBBP4
HDAC1
PSMC4
NOL9
EFTUD2
MYBBP1A
DDB1
NAP1L1
NAP1L4
RPS7
RPS11
RPL38
RPL3
RPS14
RPS4X
RPL23
RPS3A
RPL10
RRBP1
P3H1
IK
HIST1H2AC
BBX
TUBB2A
DNAJA1
MSH6
DNAJA3
CCDC47
AIFM1
GALK1
SAMHD1
CAND2
SAP30
SAP30L
SIN3A
FOXK1
ARID4A
SIN3B
SAP130
BRMS1
ARID4B
FOXK2
USP11
TUBB3
BRMS1L
SUDS3
ING2
ING1
TNRC18
BAHCC1
HIGD1A
MCM7
IPO4
HAX1
UBR5
TUBB6
B
Input: 5%
AP: Halo
Halo
Halo-Fam60A
Halo
Halo-Fam60A
FAM60A
SIN3A
RBBP4
RBBP7
HDAC1
SUDS3
SAP30
Halo
GAPDH
C
MOTIFs(M)
M1 =MOTIFΔ1 TYWKR>VFWLI
M2 =MOTIFΔ2 CCGI>MMAL
M3 =MOTIFΔ3 YKGR>FIGI
M4 =MOTIFΔ4 KPCC>LAMM
148
FAM60A spliced variant 148-221
221
MOTIF replaced with VFWLI MMAL FIGI IAMM
Human MASGSNRTPVFSFLDLTYWKRQKICCGIIYKGRFGEVLIDTHLFKPCCSNKKAAAEKPEEQGPEPLPISTQE
Sandgrouse MSAGSSRTPVFSFLDLTYWKRQKVCCGIIYKGRFGEVLIDTHLFKPCCSNKKSATEKPEQEGPQSPAVSTQE
Kingsleyae LASGSSRSPVFSFLDLTYWKRQKVCCGIIYKGRFGEVLIDPHLFKPCCRKKQQQQQQQQEEEDETQESQE
Orange clownfish LSPGTSLPPAFSFLDLTYWKRQKVCCGIVYKGRYGEVLIDPHLYKPCCQRKPQLHQPQEEGGLLEEEAAGE-
Rainbow trout LSPGATHSPVFSFLDLTYWKRQKVCCGIIYKGRFGEVLLDPHLYKPCCQKKRQHQPKEEGEEEAQASRA---
Zebra mbuna SKQRRSAPSIFSFLDRSYWKRQKVCCGIVYKGRFGEVIIDPRLFKPCCSSKKQKTLTSPTQVATHLPDT---
Asian bonytongue AKQTRSTPSLFSYLDLSYWKRQKVCCGIIYKGRFGEVMIDPRLFKPCCSSKKQEAPPPEPHP----------
Pundamilia nyererei SKQRRSAPSIFSFLDRSYWKRQKVCCGIVYKGRFGEVIIDPRLFKPCCSSKKQKTLTSPTQVATHLPDT---
Red-bellied piranha TKQRRPNPAGFSFLDLSYWKRQKVCCGIVYKGRFGEVIIDPRLFKPCCSTKERAALTSNADSQNSKSP----
Ocellaris clownfish SKRRRSTPSIFSFLDRSYWKRQKVCCGIVYKGRFGEVIIDPRLFKPCCSSKKQKTLTSTQV-ATHMPDTL--
Bicolor damselfish SKQRRSTPSIFSFLDRSYWKRQKVCCGIVYKGRFGEVIIDPRLFKPCCSSKKQKTLTSTQV-AAHMPDTL--
**:** :******:****:****:***::* :*:**** *
M1 M2 M3 M4

Fig. S4

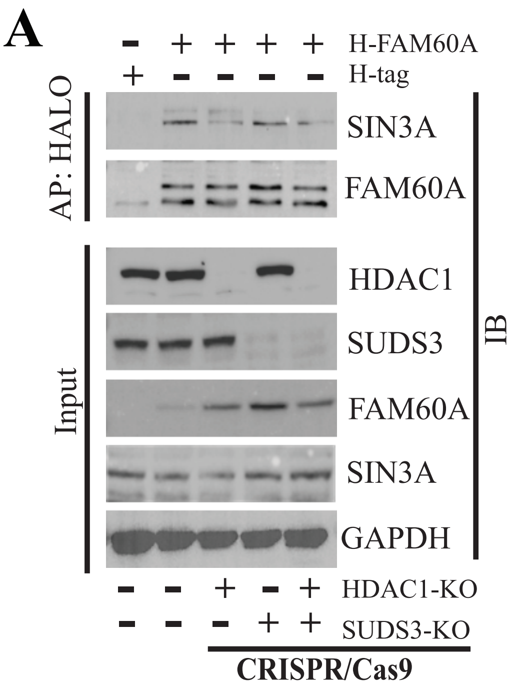
A
H-FAM60A
H-tag
AP: HALO
SIN3A
FAM60A
Input
HDAC1
SUDS3
FAM60A
SIN3A
GAPDH
IB
HDAC1-KO
SUDS3-KO
CRISPR/Cas9

**Fig. S5**

**A** **KEGG pathway analysis for Fam60A associated mRNA**

| Category | Term | RT | Genes | % | P-value | Benjamini |
|---|---|---|---|---|---|---|
| KEGG_PATHWAY | Ribosome | RT | 14 | 7.6 | 6.70E-11 | 7.30E-09 |
| KEGG_PATHWAY | Coronavirus disease - COVID-19 | RT | 15 | 8.1 | 3.40E-10 | 1.90E-08 |
| KEGG_PATHWAY | Spliceosome | RT | 8 | 4.3 | 6.50E-04 | 2.40E-02 |
| KEGG_PATHWAY | Cardiac muscle contraction | RT | 4 | 2.2 | 2.20E-02 | 6.00E-01 |
| KEGG_PATHWAY | Prion disease | RT | 6 | 3.2 | 3.80E-02 | 8.40E-01 |
| KEGG_PATHWAY | Viral carcinogenesis | RT | 5 | 2.7 | 5.10E-02 | 8.80E-01 |
| KEGG_PATHWAY | Huntington disease | RT | 6 | 3.2 | 5.80E-02 | 8.80E-01 |
| KEGG_PATHWAY | Oxidative phosphorylation | RT | 4 | 2.2 | 6.40E-02 | 8.80E-01 |
| KEGG_PATHWAY | Necroptosis | RT | 4 | 2.2 | 9.60E-02 | 1.00E+00 |

*The Benjamini-Hochberg procedure controls the false discovery rate for more effective multiple testing than the more conservative Bonferroni method. RT: Functional Related Terms

**B**

| Genes involved in Spliceosome | | |
|---|---|---|
| RNA, U5D small nuclear 1(RNU5D-1) | RG | Homo sapiens |
| RNA, U6 small nuclear 8(RNU6-8) | RG | Homo sapiens |
| RNA, variant U1 small nuclear 15(RNVU1-15) | RG | Homo sapiens |
| RNA, variant U1 small nuclear 4(RNVU1-4) | RG | Homo sapiens |
| U1 spliceosomal RNA(LOC124904613) | RG | Homo sapiens |
| U1 spliceosomal RNA(LOC124904621) | RG | Homo sapiens |
| heat shock protein family A (Hsp70) member 1A(HSPA1A) | RG | Homo sapiens |
| heterogeneous nuclear ribonucleoprotein A1(HNRNPA1) | RG | Homo sapiens |

Fig. S6

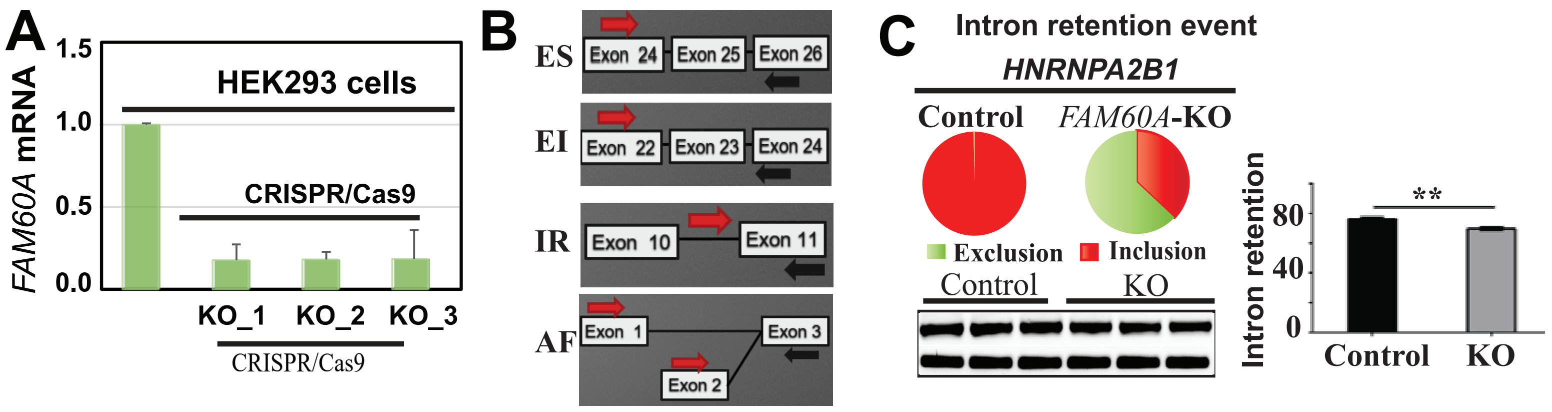
A
HEK293 cells
FAM60A mRNA
1.5
1.0
0.5
0.0
CRISPR/Cas9
KO_1
KO_2
KO_3
CRISPR/Cas9
B
ES
Exon 24
Exon 25
Exon 26
EI
Exon 22
Exon 23
Exon 24
IR
Exon 10
Exon 11
AF
Exon 1
Exon 3
Exon 2
C
Intron retention event
HNRNPA2B1
Control
FAM60A-KO
Exclusion
Inclusion
Control
KO
Intron retention
80
40
0
**
Control
KO

**Supplementary Figure legend:**

**Suppl. Fig. S1: Domain specific protein-protein interaction of the SIN3A Protein. a**, schematic-diagram illustrating various deletions in the *SIN3A* gene. **b**, the SIN3A deletion mutants were Halo tagged and transfected in to HEK 293 cells, the cells were then subjected to immunoblotting analysis using anti-SIN3A and anti- tubulin antibodies and tubulin as a loading control. **c**, Halo-tagged SIN3A deletion mutants were transfected into HEK293T cells. Halo-Tag TMRDirect fluorescent ligand (red) was used to label Halo-Tag proteins; DNA was stained with Hoechst dye (blue). **d** and **e**, Halo-SIN3A deletion mutants were ectopically expressed in HEK293T cells followed by Halo affinity purification and the purified proteins were subjected to proteomics analysis (log2FC $\geq 2$, FDR < 0.05) and visualized in a heatmap (green and black) (**d**), identified SIN3A domain specific interaction with core subunits have been visualized in a heatmap (grey indicate no interaction).

**Suppl. Fig. S2: Cross-Species similarity of FAM60A amino acid sequences. a,** the FAM60A protein sequence was analyzed using the standard protein blast tool on the NCBI blast website, aligning the query sequence with those in a specified target database. b, a schematic diagram depicts various deletions in the FAM60A gene. Halo-tagged FAM60A mutants were transfected into HEK293 cells, and post-halo affinity purification, the proteins were resolved on SDS-PAGE and visualized via silver staining.

**Suppl. Fig. S3: FAM60A employs various domains to facilitate interactions with different proteins.** a, full-length FAM60A, along with its deletion mutants FAM60A_Δ1 (1-148) and FAM60A_Δ4 (149-221, spliced variants), were Halo-tagged and transfected into HEK 293 cells. Post-transfection, Halo affinity purification was performed, and the proteins underwent proteomics analysis (log2FC $\geq 2$, FDR < 0.05), followed by visualization in an interaction network via Cytoscape. b, Halo-tagged FAM60A was transfected into HEK 293 cells, followed by Halo affinity purification. The purified proteins were analyzed through immunoblotting using various antibodies, with Halo and GAPDH serving as loading controls. c, the conserved motif in FAM60A's C-terminus (FAM60A_Δ4 149-221: spliced variant) was identified using the standard protein blast tool on the NCBI BLAST website. The conserved motif is highlighted and marked with asterisks at the bottom.

**Suppl. Fig. S4: Knockout of *HDAC1* and *SUDS3* in HEK293 Cells.** a, *HDAC1* and *SUDS3* were individually or combinatorically knocked out with CRISPR/Cas9. Subsequently, Halo (H) tag alone or Halo-FAM60A was overexpressed, and protein complexes were purified through affinity purification. These complexes were then subjected to immunoblotting analysis and probed with various antibodies.

**Suppl. Fig. S5: Pathway analysis of mRNA copurifying with FAM60A.** a. after transfecting Halo-FAM60A/FAM60A-Halo into HEK293 cells and performing affinity purification, RNA was isolated from the purified FAM60A complexes. RNA sequencing then revealed mRNAs enriched in RNA processing and splicing, with a focus on spliceosome pathways, as elaborated in box b.

**Suppl. Fig. S6: FAM60A in splicing regulation.** a, multi-guide CRISPR/Cas9 was used to knock out FAM60A in HEK 293 cells, with the knockout confirmed by qRT-PCR. b, Pie charts illustrate and quantify intron retention events, validated by RT-PCR. c, a schematic displays primer targeting regions for confirming splicing events like exon skipping (SE), exon inclusion (EI), intron retention (IR), and alternate first exon. **P $\leq 0.001$.
